# Causal network inference based on cross-validation predictability

**DOI:** 10.1101/2022.12.11.519942

**Authors:** Yuelei Zhang, Qingcui Li, Xiao Chang, Luonan Chen, Xiaoping Liu

## Abstract

Identifying causal relations or causal networks among molecules/genes, rather than just their correlations, is of great importance but challenging in biology and medical field, which is essential for unravelling molecular mechanisms of disease progression and developing effective therapies for disease treatment. However, there is still a lack of high-quality causal inference algorithms for any observed data in contrast to time series data. In this study, we developed a new causal concept for any observed data and its causal inference algorithm built on cross-validated predictability (CVP) can quantify the causal effects among molecules/genes in the whole system. The causality was extensively validated by combining a large variety of statistical simulation experiments and available benchmark data (simulated data and various real data). Combining the predicted causal network and the real benchmark network, the CVP algorithm demonstrates high accuracy and strong robustness in comparison with the mainstream algorithms. In particular, the CVP algorithm is robust in identifying reliable driver genes and network biomarkers from the perspective of network biology, with the prediction results outperforming the mainstream conventional methods for predicting driver genes. CRISPR-Cas9 knockdown experiments in the liver cancer have validated that the functional driver genes identified by the CVP algorithm effectively inhibit the growth and colony formation of liver cancer cells. By knockdown experiments, we demonstrated the accuracy and significance of the causality predicted by CVP and identified the novel regulatory targets of functional driver genes SNRNP200 and RALGAPB in the liver cancer. These inferred causal networks explain regulatory patterns in different biological contexts in a clear sense and provide biological insights into molecular mechanisms of disease progression from a causality perspective.

## Introduction

Causal inference from a observed data is a core problem in various research disciplines of natural science and engineering, such as biology, earth science, economics, medicine, neuroscience and machine learning. Molecular networks are the essential issue of biological systems in view of causal inference (*1*) and building high-quality molecular networks based on the observed/measured data has always been a vital problem of computational biology (*2, 3*). Effective identification of causal relations in a complex biological system can better reveal regulatory mechanisms and explain biological functions, such as gene regulations, signaling processes, metabolic pathways and disease progression (*4–6*). In particular, deriving a specific disease causal/regulatory network can reveal the basic mechanism of molecular effects to provide quantitative studies for further precise treatment.

Causal effects between molecules can often be represented by causal diagrams, with nodes representing different molecules and directed edges characterizing the direct causality between molecules(*7*). Existing mainstream algorithms for monitoring causality include the well-known Granger causality (GC) (*8, 9*), the convergent cross-mapping algorithm (CCM) (*10, 11*), transfer entropy (TE)(*12, 13*) and Bayesian theory (*14, 15*). Granger causality (GC) inference as a representative method, that is based on time series data to infer the potential causality, was proposed in 1969 (*8*), and since then GC-based methods have been widely used in many fields. Specifically, GC is based on the information of time lags or time series data, and explains the current state to be affected by past information. TE as a nonlinear version of GC method considers the asymmetry of information in time series to determine causality and is commonly used in biological systems such as neuroscience (*16, 17*) and physiology (*18, 19*). Both GC and TE are measured at the original state space. On the other hand, CCM(*11, 20*) as a complement to GC theory is measured in the delay embedding space, which is also based on time series data and reflects causal relationship between two variables (*21, 22*). All of the above algorithms measure causality with a requirement of time dependent data or time-series data, but most of the biological data are not based on time series data, such as phenotypes, stages or phases, thus unsuitable for applying such methods. In contrast, Bayesian network and structural causal model (SCM) (*23–25*) are able to handle time independent data based on statistical independence and intervention (e.g. do-calculus operation), and can identify the directed causal relations between molecules. However, these methods depend on the structure of directed acyclic graph (without loop structure) to infer causality, which limits to the application to biomolecular networks commonly with feedback loops or ring-like interactions (*26*). Therefore, a high-quality inference method of causality in real biological/molecular systems without time or structural limitation, rather than relying on time dependent data, remains an open question.

In this study, we proposed a new causal concept and the method termed cross-validation predictability (CVP), which is a data-driven model-free algorithm for any an observed data. The CVP method quantifies causal effect by cross-testing predictability and statistical test of the observed variables on any observed data. The CVP method was statistically tested with a large number of causal simulation experiments as well as fully performing validation on simulated data from existing benchmarks. Moreover, the method is also shown the superior performance and significant advance in a variety of real systems, including gene regulatory networks and other causal networks with feedback loops, by extensive comparisons with various existing methods. In particular, CRISPR-Cas9 knockdown experiments in the liver cancer have validated that the functional driver genes identified by the CVP algorithm effectively inhibit the growth and colony formation of liver cancer cells. By knockdown experiments, we demonstrated the accuracy and significance of the causality predicted by CVP and identified the novel regulatory targets of functional driver genes SNRNP200 and RALGAPB in the liver cancer. In summary, CVP is a general-purpose method for causal inference based on the cross-validation testing on predictability from any observed/measured data, and able to infer causality among the variables in an accuracy and robust manner.

## Results

### Causality detection based on cross-validation predictability

The causality from CVP method is a statistical concept based on cross-validation prediction of observed data. Assuming two variables *X* and *Y* are observed in *m* samples, we consider that *X* causes *Y* if the prediction of the values of *Y* is improved by including values of *X* in the sense of cross-validation (Fig. 1). Specifically, we assume that a variable set {*X*, *Y, Z*_1_, *Z*_2_, ⋯, *Z*_*n*–2_} includes *n* variables in the *m* samples, where 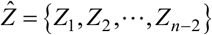. In other words, *X* and *Y* are any two variables among all *n* observed variables. All *m* samples are randomly divided into a training group and a testing group, for the purpose of cross-validation, e.g., *k*-fold cross-validation. To test causal relation from *X* to *Y*, we formally define CVP causality framework, i.e. construct two contradictory models H_0_ (null hypothesis without causality) and H_1_ (alternative hypothesis with causality) by the same *k*-fold cross-validation and further define causal strength by the difference between H_1_ and H_0_ for quantifying CVP causality as follows.

- H_0_:

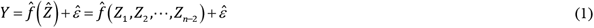 Train the regression 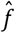 by the training group samples, and then test 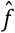 by the testing group samples in a *k*-fold cross-validation manner. We have the error 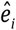 of Eqn. (1) in the *i*-th cross-validation test by the testing group samples. The total squared testing error is 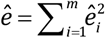 for all *k*-fold cross-validation tests.
- H_1_:

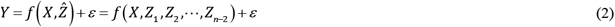 Train the regression *f* by the training group samples, and then test *f* by the testing group samples in a *k*-fold cross-validation manner. We have the error *e_i_* of Eqn. (2) in the *i*-th cross-validation test by the testing group samples. The total squared testing error is 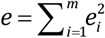 for all *k*-fold cross-validation tests. If 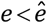, then H_1_ holds, i.e. causal relation from *X* to *Y*. And if 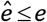, then H_0_ holds, i.e. no causal relation from *X* to *Y*.
- Causal strength (CS):

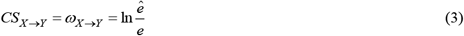

**Fig. 1.**
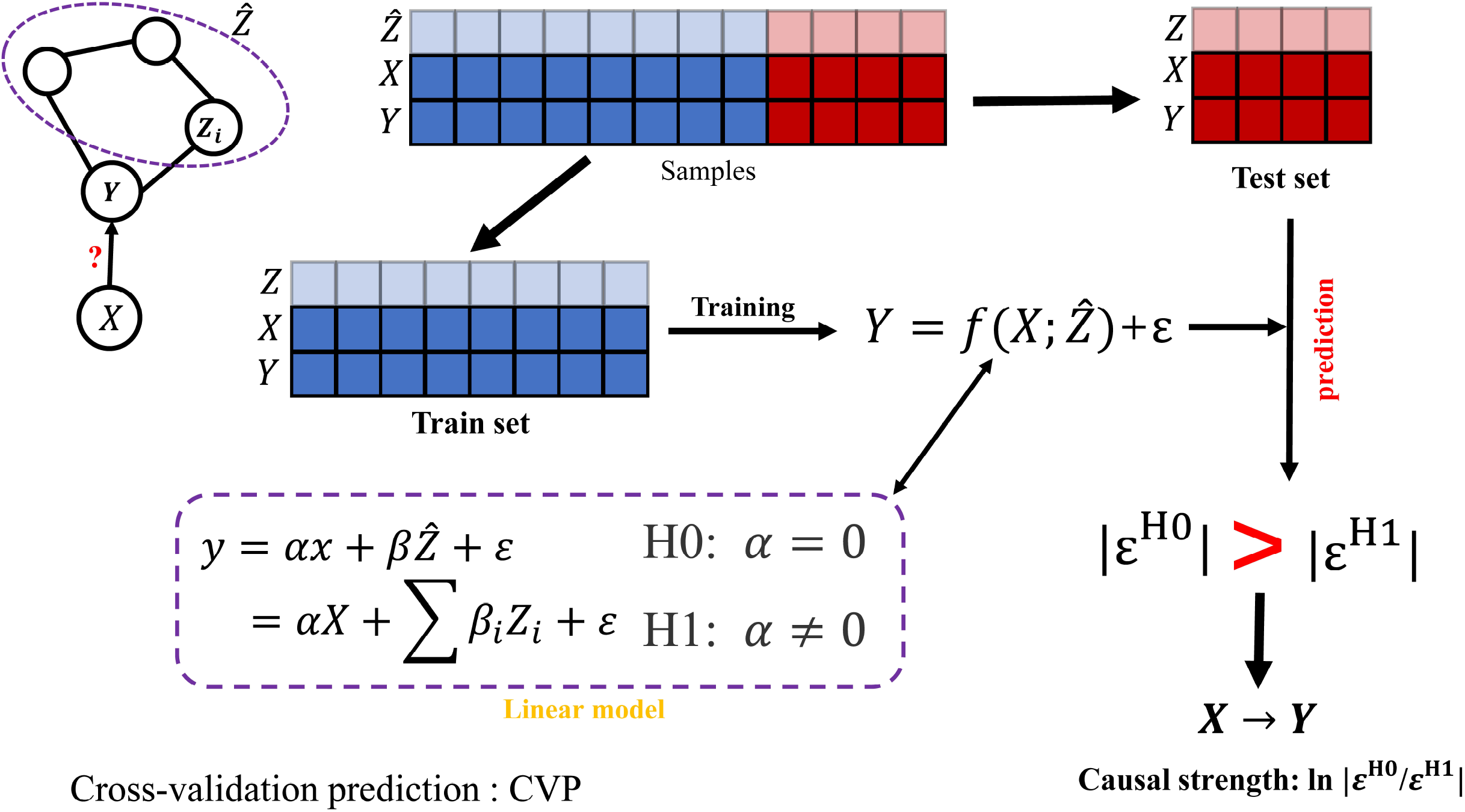
The causal prediction method based on modelling and prediction. The flowchart To infer/judge the causality from variable *X* to *Y*, all samples are divided into train set and test set based on cross-validation, and then modelling the causality equation for variable *Y* with and without *X* by train set. The test set was used to predict variable y and calculate the error *ε*.

Causal strength is the difference/distance measured between H_0_ and H_1_, i.e. 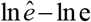. Of course, other statistical test (e.g., a paired Student’s *t*-test) can be also used to test the difference between *e* and 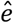, i.e. significance test.

Here, *f* is the regression equation for *Y* fitting *X* and 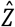, and 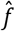 is the regression equation for *Y* fitting 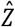 without *X*. In this work, we use the linear regression for both *f* and 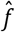. The *ε* and 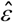 are the error terms of Eqns. (2) and (1), respectively. The error is defined as the difference between the predicted value and the true value in testing group.

The Eqns. (1) and (2) are first fitted from the training group samples, and then are tested by the testing group samples for determining the testing/predicting errors, which are summarized as 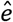 and *e* including the error of every sample in the testing group by Eqns. (1) and (2). We also ensure those errors are statistically independent of 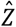 and *X* by applying appropriate the regression algorithm, e.g., least-squares regression algorithm. Thus, causal criterion of CVP is essentially based on model predictability and statistical independence.

Clearly, if H_1_ or *e* is significantly less than 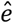 in the testing group, it means that Eqn. (2) is a better regression equation than Eqn. (1), i.e. *X* causes *Y* (Fig. 1 and Supplementary Note 1). Otherwise, H_0_ holds, i.e. *X* does not cause *Y* (Fig. 1 and Supplementary Note 1). In this paper, we also introduce causal strength (Eqn. (3)) to test the difference between *e* and 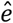. Here, since the variable set 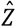 can be considered as other factors to affect variable *Y* except *X*, hence this method considers or infers the direct causality from *X* to *Y* under all other factors fixed.

### Benchmarking dataset

The superiority of the CVP algorithm was validated by different types of data, including the DREAM (Dialogue on Reverse Engineering Assessments and Methods, https://dreamchallenges.org) challenges (especially DREAM3 and DREAM4) and biosynthesis network (IRMA data) from Saccharomyces cerevisiae (*27*), and further a variety of real data, such as the SOS DNA repair network in Escherichia coli(*28, 29*), yeast data (DDGni Yeast) (*30*), the human HeLa data(*31*), the TCGA cancer data (https://portal.gdc.cancer.gov/) and the *E. coli* data from GEO database(*32*). The DREAM challenges and IRMA data demonstrate the adaptability of external disturbances to the CVP algorithm. For each dataset, the performance of the CVP algorithm was evaluated against eight of the most popular algorithms, i.e. Nonlinear ODEs (*33*), GENIMS (*34*), PLSNET (*35*), GENIE3_ET_all, GENIE3_ET_sqrt, GENIE3_RF_all and GENIE3_RF_sqrt (*36*), and TIGRESS (*37*). In this study, we used the default parameters of the algorithm in order to make a fair comparison with those methods (Supplementary Note 2). Moreover, the DREAM challenges dataset was used for extensive benchmarking, and the algorithms that performed best in the DREAM challenges were included in the comparison results.

### Causal effect estimation for simulated data

Two simulated causal networks were used to generate simulation data and test the effectiveness of the CVP algorithm. One network includes three nodes with different causal links (Fig. S1, Fig. S2a-S2e), which produces corresponding simulation data (Supplementary Note 3 and Tab. S1). The CVP algorithm was used to quantify the causality from the simulated data (Fig. S2f-S2j). The results show that the CVP algorithm can accurately measure the causal network structure among the three nodes from the simulated data with few false links or interactions (Supplementary Note 3 and Fig. S2f-S2j). The CVP algorithm not only can identify the true causal relations but also eliminate the false causal links, i.e. indirect causal links caused by cascade or confounding factors (Fan-out) or collision factors (Fan-in) (Supplementary Note 3 and Fig. S2).

Other nine causal networks were constructed from a basic regulatory network including 11 nodes with one center node and ten neighbor nodes and the center node was regulated by all the neighbor nodes, and then the neighbor nodes were removed from the initial regulatory network one by one to form nine structures or cases (Fig. S3a). Each structure or case was used to produce the corresponding simulation data of 11 variables (Tab. S2). The CVP algorithm was also used to infer the causality between the center node and ten neighbors based on the simulation data of 11 variables (Supplementary Note 4 and Fig. S3b-S3c). The results show that the CVP algorithm can accurately infer the network structure and causality among the 11 nodes from the simulated data with an accuracy rate to almost 100% for both existent and non-existent edges in the 9 cases (Supplementary Note 4 and Fig. S3b-c).

### Causal effect estimation for data in DREAM challenges

The DREAM challenges (*3, 38*) have been widely used as a benchmark dataset for causal inference. We obtained 4 datasets from 4 networks with 10 genes/nodes separately from DREAM3 and DREAM4, and the 4 networks were marked Network1~Network4 (Fig. 2a-2d) and Supplementary Note 5).

**Fig. 2.**
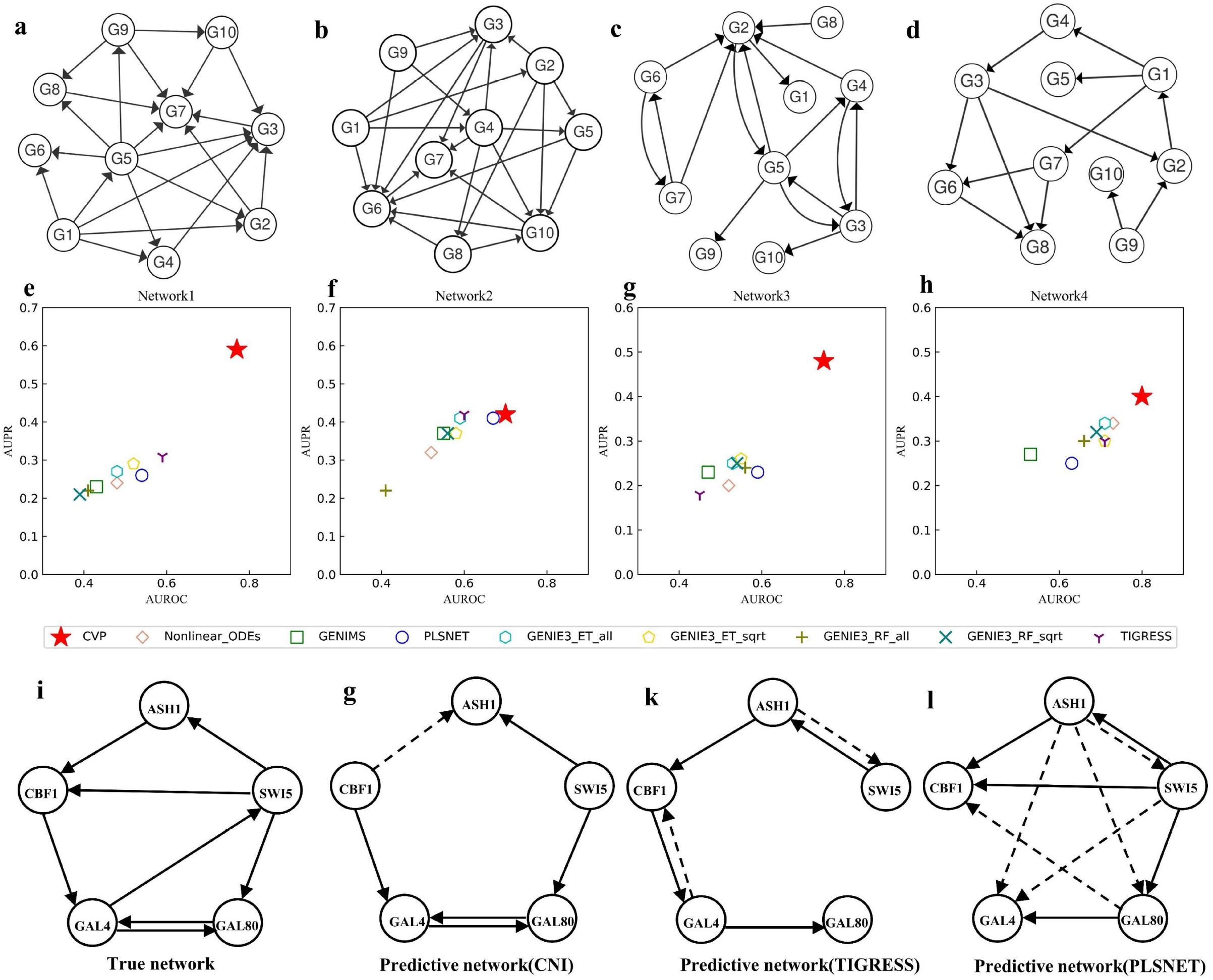
The performance of different algorithms on DREAM challenges and IRMA datasets. a,b,c,d are the real networks from Network 1, Network 2, Network 3, and Network 4 from the DREAM challenges. e,f,g,h are the performance of the CVP and other eight methods for inferring Network 1, 2, 3 and 4 respectively. The horizontal coordinates are the AUPR values and the vertical coordinates are the AUROC values. The shape of the dots indicates the different algorithms, with dots closer to the top right corner indicating better algorithm performance. i,g,k,l show the network for IRMA data. i shows the ground truth network, and g and k, and l are the inferred networks of the CVP, TIGRESS, and PLSNET algorithms, respectively. The solid lines represent the accurate edges (true positive edges) to be inferred by the methods, and the dashed lines represent the wrong edges (false positive edges) to be inferred by the methods for g, k and l.

We compared the performance of the CVP algorithm with other eight algorithms, such as Nonlinear ODEs, GENIMS, PLSNET, GENIE3_ET_all, GENIE3_ET_sqrt, GENIE3_RF_all, GENIE3_RF_sqrt and TIGRESS based on two datasets of two networks in DREAM3 challenge (Fig. 2a and 2b), that one is a non-time series data for Network1 and another is a time series data for Network2 (Supplementary Note 5). Clearly, the results demonstrated that the CVP algorithm performed better than other algorithms for the two networks (Fig. 2e and 2f). The closer the position locates on the top right of the coordinate system, the better performance of the algorithm is (Fig. 2e-2h). For the Network1 (Fig. 2a), we can clearly find that CVP algorithm is much higher than other algorithms in AUROC (area under the receiver operating characteristic curve) and AUPR (area under the precision-recall curve)) values, with AUROC value reaching 0.77 and AUPR value 0.59 (Fig. 2e, Fig. S4a and S4e). TIGRESS ranked second among all the algorithms for Network1, with an AUROC of 0.59 and an AUPR of 0.31, which are much lower than the CVP algorithm (Fig. 2e, Fig. S5a and S5e). For the Network2 (Fig. 2b), the CVP and PLSNET algorithms are close in performance, whose AUROC values are separately 0.7 and 0.67, and the AUPR values are separately 0.42 and 0.41, respectively (Fig. 2f, Fig. S4b and S4f). The CVP algorithm slightly outperforms PLSNET algorithm and performs best (Fig. 2f).

Similar to the DREAM3 networks, DREAM4 also includes simulation data and corresponding real networks. We chose two datasets of two networks from DREAM4 challenge (Fig. 2e and 2d), and one is a non-time series data for Network3 and another is a time series data for Network4 (Supplementary Note 5). The CVP algorithm outperformed the other algorithms for the two datasets by a significant margin, with AUROC values of 0.75 and 0.8 in Networks3 and Network4 (Fig. 2g, 2h and Fig. S4c, S5d), respectively. In Network 3, all AUROC values of other algorithms are lower than 0.6, and AUPR values vary around 0.25 in similar performance. It may be because the complexity of the network leads to the unsatisfactory inference of the algorithm, while the AUPR value of CVP is 0.48, which still maintains the best performance (Fig. 2g). The GENIE3_ET_all, GENIE3_ET_sqrt, GENIE3_RF_sqrt, Nonlinear ODEs and TIGRESS algorithms performed well, with AUROC value of approximately 0.7 for Network4 (Fig. 2h and Fig. S4d), but the CVP algorithm performed as high as 0.8 (Fig. 2h and Fig. S4d), which still maintained the best performance.

For the DREAM challenges dataset, we totally tested 40 datasets at different scales (Supplementary Note 5 and Fig. S4-S7), and the CVP method performed better than other algorithms (Supplementary Note 5 and Fig. S4-S7). For the network with size 10, the CVP algorithm always has far better AUROC and AUPR values than the other algorithms (Fig. 2e-2h, Fig. S4 and Fig. S5). We also compared the accuracy of the algorithm for networks with size 50 and 100, and the CVP algorithm outperformed the other algorithms by a wide margin in 30 datasets, reflecting the accuracy and stability of CVP in synthetic datasets (Supplementary Note 5 and Fig. S4-S7).

### Causality detection in synthetic datasets

The IRMA network is a synthetic network (*27*) embedded in Saccharomyces cerevisiae, cultured in vivo to obtain ground biological data, and is widely used as benchmark data in inferential modelling. This network contains five genes and eight regulatory edges, whose structure is known (Fig. 2i).

The IRMA network is relatively small, and the values of AUROC and AUPR are relatively sensitive to small changes of the threshold in the inferred results. Obviously, the CVP algorithm is not subject to this limitation, and the AUROC and AUPR values are the highest compared with the other algorithms, which are 0.79 and 0.75 (Fig. S8), respectively. For the eight edges in the ground truth network (Fig. 2i), the CVP algorithm accurately inferred five edges, with only one false positive edge (Fig. 2g). Relatively good algorithms, such as TIGRESS inferred four edges and two false positive edges (Fig. 2k and Supplementary Note 6), or PLSNET inferred five edges and five false positive edges (Fig. 2l). However, these algorithms only provide the intensity of regulation without a specific threshold, which makes it difficult to determine the optimal subnet. The CVP algorithm not only yields the optimal directed network, but also quantifies the causal strength (Fig. S8d). In the latest study, the best algorithm of this network inference is the BINGO algorithm (*7*) with five inferred edges and one false positive, in agreement with our accuracy, but the method is restricted to the data based on time series data. Therefore, the CVP algorithm has a high performance in in vivo synthesis networks.

### Reconstruction of real gene regulatory networks

A real network is usually a complex nonlinear system, whose causal structure is generally difficult to be inferred. We focused on gene regulatory networks in biological systems, and applied the CVP algorithm to five real datasets of gene regulatory networks, i.e. the SOS DNA repair network data, Saccharomyces cerevisiae cell cycle data, human HeLa cell cycle data from literatures and BLCA dataset from TCGA. The performance of different algorithms was estimated using the AUROC or AUPR for each gene regulatory network.

#### The SOS DNA repair network

The SOS DNA repair network is the most commonly benchmark data set, which is the real non-time series data verified by experiments in *E. coli* (*28, 29*). It is a non-linear complex network consisting of 9 genes and 24 edges(*28, 29*) (Fig. 3i and Supplementary Note 7). By comparing the inference results based on the real data, the CVP algorithm outperformed the other algorithms for the SOS network (Fig. 3a, 3e and 3j). The AUROC value of the second highest ranking algorithm is 0.51, and the AUROC value of the CVP algorithm is 0.75, which is at least 24% higher than other algorithms (Fig. 3a and Fig. S9a). The AUPR of the CVP is 0.63, and the second AUPR in the other algorithms is Nonlinear_ODEs and PLSNET with 0.35, that is 28% lower than the CVP (Fig. 3e and Fig. S9b). At the same time, high false positives are always the main problem facing inferred GRNs (gene regulatory networks), while the CVP algorithm infers 16 edges in the SOS network with only 3 false positives (Fig. 3j), while other methods infered the optimal network with at least half of the false positive edges (Fig. 3k and 3l). It indicates that our algorithm introduces fewer redundant edges and is able to accurately infer the true network (the accuracy of CVP is 81%) (Fig. 3j, Supplementary Note 7 and Tab. S5).

**Fig. 3.**
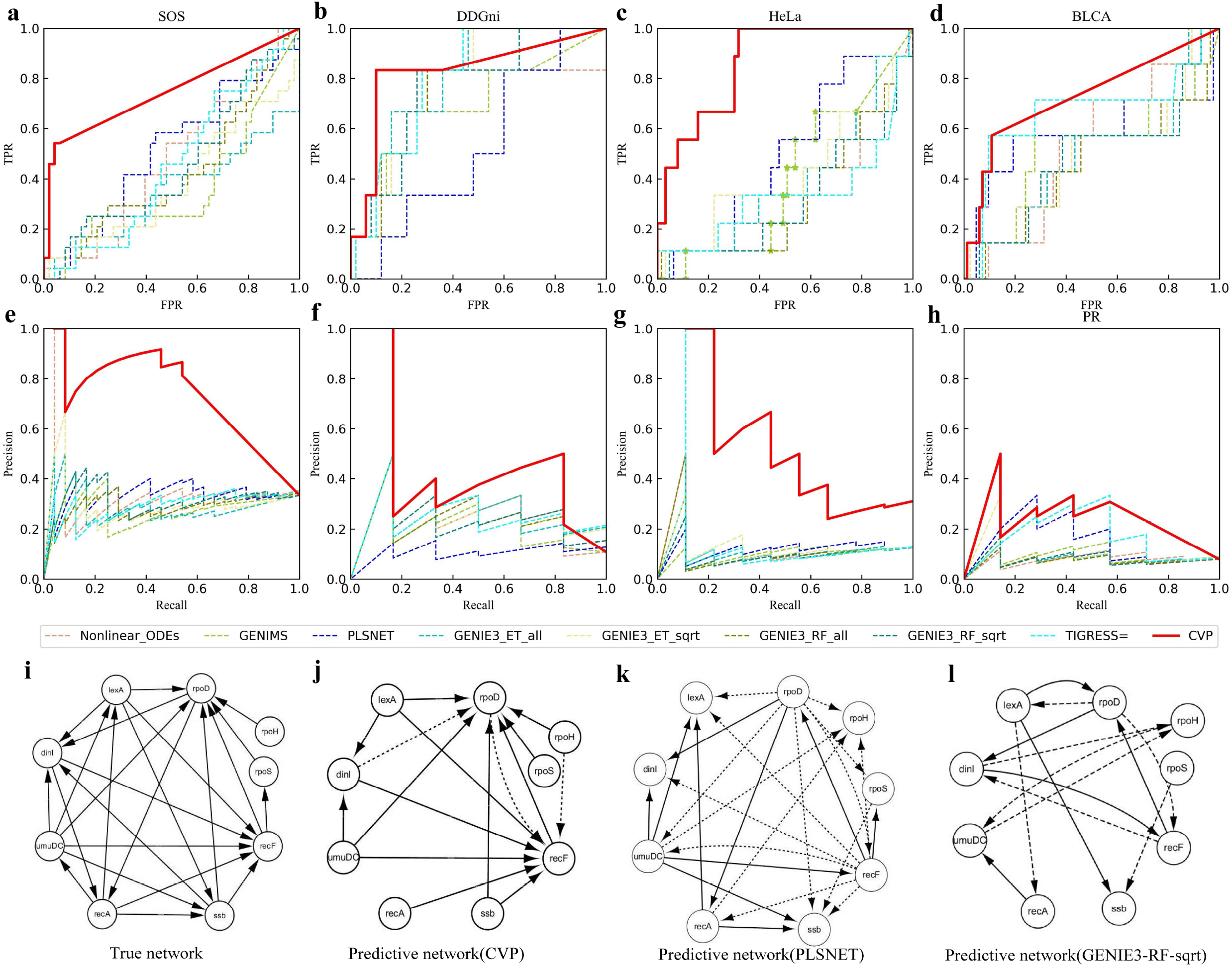
The performance of different algorithms on the different real dataset. a-h show the performance of the nine algorithm s (Nonlinear ODEs, GENIMS, PLSNET, GENIE3_ET_all, GENIE3_ET_sqrt, GENIE3_RF_all, GENIE3_RF_sqrt, TIGRESS and C VP) on the SOS, DDGni, HeLa, and BLCA datasets respectively. a,b,c,d show the ROC curves for the SOS, DDGni, HeLa, and BLCA datasets, and e,f,g and h show the PR curves for the SOS, DDGni, HeLa, and BLCA datasets, where the different colore d lines indicate different algorithms. **i** shows the real network for SOS dataset. j, k and l denote the inferred networks from CV P, PLSNET GENIE3-RF-sqrt algorithms for the SOS data, respectively. The solid lines represent the accurate edges (true positive edges) to be inferred by the methods, and the dashed lines represent the wrong edges (false positive edges) to be inferred by t he methods for j, k and l.

#### The yeast cell cycle data

This data was obtained by the authors of DDGni(*39*) from the accession number GSE8799 on the Gene Expression Omnibus (GEO, http://www.ncbi.nlm.nih.gov/geo/) database. The DDGni data was with 8 genes, 6 edges, and contains 30 time points of gene expression data from case of yeast cell cycle dataset. The time series data and golden standard regulatory network was reported in literature(*39*), and this data is also widely used as a benchmark for assessing the inference of GRNs (*33*). We plotted the ROC (Fig. 3b) and PR (Fig. 3f) curves for CVP and the other eight algorithms. Overall, each algorithm performed well for this network, but the CVP algorithm was still the highest in AUROC and AUPR values (Fig. 3b and 3f). The AUROC value of the CVP algorithm is 0.83 (Fig. S9a), that for the other algorithms are basically between 0.7 and 0.8, and only PLSNET has a value of 0.53 (Fig. 3b and Fig. S9a), probably due to a low AUROC due to high false positives. Similarly, the CVP algorithm has an AUPR of 0.47 (Fig. S9b), while the other eight algorithms have an AUPR of no more than 0.4, with the highest being the GENIE3-RF-sqrt algorithm at 0.38(Fig.3b and Fig. S9b). Therefore, our algorithm is more robust and accurate to infer the real network of yeast cell cycle data.

#### The human HeLa cell cycle dataset

To compare the performance of the CVP algorithm with other algorithms, we used a biological experiment data, namely the HeLa cell cycle gene expression data (*31*). Sambo et al.(*40*) reported a subnetwork with corresponding time series data, which is then often used as a benchmark (*41–43*) for evaluating algorithms to construct GRNs. The expression data of the ground truth network with nine regulatory relationships among nine genes was used to evaluate the performance of the CVP and other algorithms. The ROC and PR curves of all algorithms were shown in Fig. 3c and 3g, and it is clearly that the curve of the CVP algorithm is always much higher than others (Fig. 3c and 3g). The AUROC value of the CVP algorithm is 0.86 (Fig. S9a), while other algorithms are close to 0.5, which is not higher than random enumeration (Fig. 3c and Fig. S9a). At the same time, the AUPR values of the all nine algorithms (CVP, Nonliner_ODEs, GENIMS, PLSNET, GENIE3_ET_all, GENIE3_ET_sqrt, GENIE3_RF_all, GENIE3_RF_sqrt and TIGRESS) are 0.56, 0.15, 0.10, 0.14, 0.13, 0.15, 0.14, 0.12 and 0.21 (Fig. 3g and Fig. S9b), respectively. And we can see that the CVP is also better than other eight algorithms in AUPR values. When inferring the network, the CVP algorithm identified the largest number of true edges while ensuring the least number of false positives (Fig. S10). Thus, the results showed that the CVP algorithm outperformed other algorithms for different indexes and the inferred results of the CVP algorithm were closer to the true network than other algorithms for the human HeLa cell cycle dataset (Fig. 3c, 3g, Fig. S9 and S10).

#### The BLCA dataset in TCGA

The occurrence and development of cancer involve complex molecular regulation, monitoring regulation mechanism among genes can be beneficial to prevention and cure of different cancer efficiently. Here, we took BLCA (bladder urothelial carcinoma) cancer as an example to analyse the GRNs of oncogenes. The BLCA data is the RNA-Seq data from TCGA (the cancer genome atlas) database, but there is no ground truth network for this data. We extracted the corresponding bladder cancer pathway (hsa05219) from KEGG (Kyoto Encyclopedia of Genes and Genomes) database (https://www.genome.jp/kegg/) and obtained a local network as the golden standard regulatory network for BLCA data. And the CVP algorithm inferred a network with an accuracy of 87%, indicating that our algorithm constructed a network very close to the golden standard regulatory network. The CVP algorithm was also performed best among other algorithms (Fig. 3d and 3h). The highest AUROC value for the other algorithms was 0.66, and the CVP reached an AUROC value of 0.73 (Fig. 3d and Fig. S9a). The highest AUPR value for the other algorithms was 0.20, and the CVP reached an AUPR value of 0.24 (Fig. 3h and Fig. S9b). We also compared the results of other four cancer datasets, UCEC, LIHC, PAAD and STAD (Supplementary Note 8 and Fig. S11), and the CVP algorithm still performed best in the AUROC index. Although the AUROC value was slightly lower relative to the other real data sets, the main reason was stemmed from the “false positives”. Due to the incomplete regulation relationship in KEGG pathways, the “false positives” in here are not necessarily real false positives by these algorithms. It is possible that some “false positives” edges are real regulations but not in the KEGG pathways, so they may be new regulatory edges in tumors(*44*).

#### A large-scale network

For validating the application scope of the CVP algorithm, we compared the performance of the CVP algorithm and other five algorithms on the data of a large-scale network, and the GSE20305(*32*) dataset was used to provide a real time series data for *E. coli* under cold stress conditions from GEO database. We obtained the standard network by integrating DREAM5 with the *E. coli* data and RegulonDB database (version 9.0) (*45*) to form the final benchmark network containing 1484 genes and 3080 edges(*46*). We compared the accuracy of the CVP algorithm and other five algorithms (Nonlinear ODEs, GENIMS, PLSNET, GENIE3_RF_sqrt and TIGRESS) to infer the GRN for this network and the CVP algorithm still maintained the highest accuracy (Fig. S12). For the large-scale network, the accuracy of the CVP algorithm is 0.89 (Fig. S12), and it is better at least 20% than other five algorithms (Fig. S12). It means that the inferred network from the CVP algorithm is closer to the true network than the networks from other algorithms (Fig. S12) even if to a large-scale network.

All in all, the CVP algorithm recovered different causal networks from the real data (benchmark), and has consistently outperformed other advanced algorithms significantly.

### Detecting causal networks in multiple contexts

The CVP algorithm was also considered to infer causal networks for other four systems, a planktonic food chain network, a causal network for the number of people treated for air pollution and respiratory diseases in Hong Kong, and two networks for the spread of new coronavirus infections in two regions of Japan.

#### The plankton food chain

The first example was from a time-series data on species abundance of plankton isolated from the Baltic Sea, where the food web was sampled and tested twice a week for over 2300 days(*47–49*). The network was constructed by the five planktonic species cyclopoids, protozoa, rotifers, picocyanobacteria and nanophytoplankton with six regulations (Fig. 4a). The CVP algorithm inferred seven causal edges (Fig. 4b) that are basically consistent with the ground truth network, and these edges point in a way that is consistent with the biological laws of the food chain (Fig. 4a-4b). It is worth noting that our algorithm inferred two false positive edges, i.e., nanophytoplankton to cyclopoids and picocyanobacteria to cyclopoids, and the two edges did not exist in the ground truth network (Fig. 4a-4b).

**Fig. 4.**
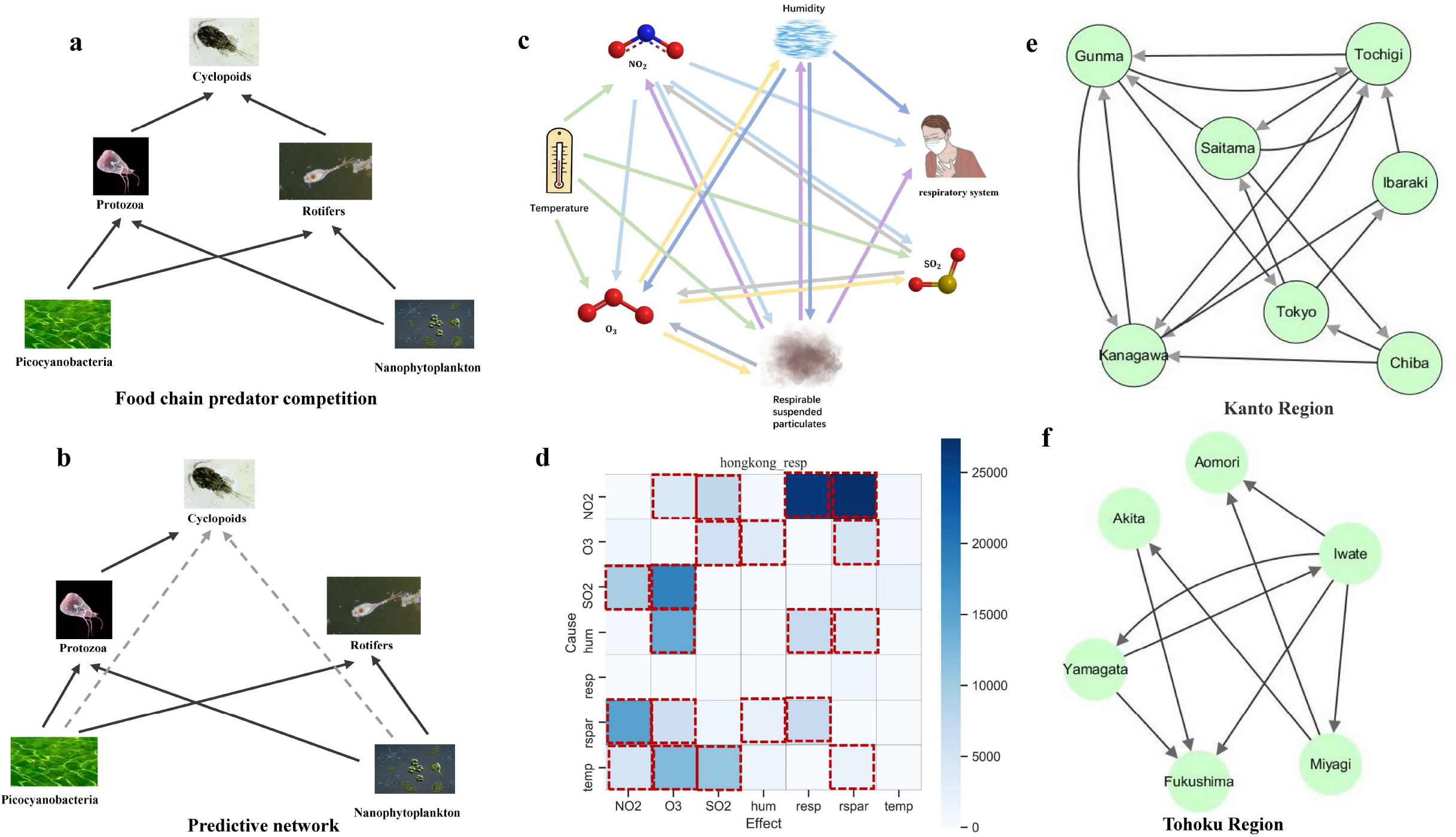
Detecting causal networks in real biological systems. a. A food chain network of plankton. b. Causality of plankton de tected by the CVP algorithm. c. The detected the causal network for respiratory diseases by the CVP algorithm. d. The inferred impact of environmental interactions on respiratory diseases by the CVP algorithm with darker heat map colors indicating stronger causal relationships. e and f show the reconstructed the network of the number of people spreading New Coronavirus by the C VP algorithm in Kanto and Tohoku of Japan, respectively.

The regulation edge from nanophytoplankton to cyclopoids was inferred by the CVP algorithm (Fig. 4b), and this regulation did not exist in the ground truth network (Fig. 4a). However, this predation or intake relationship for cyclopoids ingesting nanophytoplankton has been proved by a recent research work (*50*), and it means that the causality from nanophytoplankton to cyclopoids is true by the CVP algorithm for the plankton food chain (Fig. 4b).

Another regulation edge from picocyanobacteria to cyclopoids was also inferred by the CVP algorithm (Fig. 4b), and this regulation was not marked in the ground truth network (Fig. 4a). A research work found that cyclopoids can take cyanobacteria as food (*51*), and the picocyanobacteria is a kind of cyanobacteria with the smallest cell-size. So, the causality from picocyanobacteria to cyclopoids inferred by the CVP algorithm may be true for the food chain of the five planktonic species (Fig. 4b).

From the above results, we can see that the existent plankton food chain of the five planktonic species (Fig. 4a) is incomplete, and the CVP algorithm can be used to supplement the potential causality for the plankton food chain among the five planktonic species (Fig. 4b).

#### The causal network of respiratory inpatients and air pollution

The second example we considered is from Hong Kong air pollution data and the data of respiratory patients collected from major hospitals of Hong Kong from 1994 to 1997 (*52–54*). The CVP algorithm was used to construct the causal network among NO_2_, temperature, humidity, O_3_, respirable suspended particulates, SO_2_ and respiratory system (Fig. 4c and 4d). The CVP algorithm identified humidity, NO_2_ and respirable suspended particulates as the main causes of respiratory disease (Fig. 4c and 4d), which is consistent with prior research (*55–57*). The CVP algorithm found bidirectional causality for NO_2_ and respirable suspended particulates, and similar results were found in existing studies(*58*). Moreover, this is also consistent with the causality inferred by the prior method, e.g. PCM algorithm(*5*). The true causality between the pollutants SO_2_ and O_3_ was reported to be bidirectional(*59, 60*), which is the same as our results (Fig. 4c and 4d), but the PCM algorithm only monitored unidirectional causality from SO_2_ to O_3_ (*5*). Notably, the CVP algorithm revealed unidirectional causality for NO_2_ and O_3_, strongly related to the ability of NO_2_ in destroying O_3_, and similar discoveries are also reported in the currently available literature(*61, 62*). Here, we can find that the CVP algorithm is able to identify a causal network for the six pollutants on the respiratory system, including both unidirectional and bidirectional causation. Thus, the CVP algorithm screens for adverse factors for respiratory disease and then effectively assesses the relationship between air pollution and health.

#### New coronavirus transmission network in the region of Japan

COVID-19 is a highly infectious disease capable of causing mild or severe infection and even death in humans(*63*). Therefore, it is crucial to forecast the transmission areas of new coronaviruses. Only by obtaining accurate transmission rules and then adjusting the prevention and control measures in time, the transmission area and scale can be effectively controlled. The CVP algorithm is a data-driven approach to build causal networks across regions of transmission based on the number of infected people in different regions, and indirectly infer the laws of infection.

Here, data of daily new COVID-19 cases were collected for 13 prefectures in the Kanto and Tohoku regions of Japan from January 15, 2020 to December 13, 2020 (Supplementary Note 11). We inferred the spread network of the epidemic by the CVP algorithm in the Kanto (Fig. 4e) and Tohoku (Fig. 4f) regions of Japan (*64*). As an international transportation hub, the Kanto region is also a key area for monitoring the situation of the novel coronavirus outbreak in Japan. Taking the economic hub of Tokyo as an example, it spread to Saitama and Ibaraki and cascaded to other regions (Fig. 4e). The Narita international airport, which locates in Chiba, is a major airport to handles international passengers for Tokyo. And many infected international passengers go to Tokyo from Narita international airport. Hence, it is reasonable that the transmission from Chiba to Tokyo in Kanto region (Fig. 4e). Thereafter, the number of infected people in various regions of Japan has an outbreak of the epidemic.

Compared to the Kanto region, the epidemic in the Tohoku region is relatively mild and the transmission network is relatively sparse, which is inextricably linked to the geographical location and mobility of the population in the Tohoku region. For example, Iwate, where the COVID-19 was last seen, has a low population density, few foreign visitors and a small transient population, and at the same time began early to study countermeasures to strictly prohibit the inflow of guests from outside the prefecture in order to prevent infection by the new coronavirus. This is in good agreement with our inference that Iwate shows a tendency towards more external transmission, mainly due to its strict control policy to avoid internal transmission as much as possible.

### CVP reveals the gene regulations and functional genes during cancer progression

Presently, gastric and lung cancers are the major cancers that threaten human life(*65*). Identifying genes and regulation in cancer progression will not only improve our understanding of the biology of the process, but also provide new targets for diagnosis and treatment. We got expression profiles for gastric cancer and lung cancer in TCGA database.

In this study, we used CVP to infer the regulatory network for early (Stage I and II) and late gastric cancer (Stage IV) and identified 15 out-degree hub genes involved in the regulation network of late gastric cancer as the functional genes in cancer progression (Fig. 5c and Supplementary Note 9). Notably, it has been demonstrated through literatures, the 14 of 15 functional genes have been reported as association genes of gastric cancer and play a crucial role in progression of gastric cancer (Supplementary Note 9). By comparing the local regulatory network of the 15 functional genes in early and late gastric cancer (Fig. 5a-5b), we found that there is a similar network structure for the 15 functional genes from early (Fig. 5a) with average out-degree 9 to late gastric cancer (Fig. 5b) with average out-degree 10. It is consistent with the reported genes, e.g., VEGFA was reported to take part in tumor growth and metastasis in gastric cancer(*66, 67*).

**Fig. 5.**
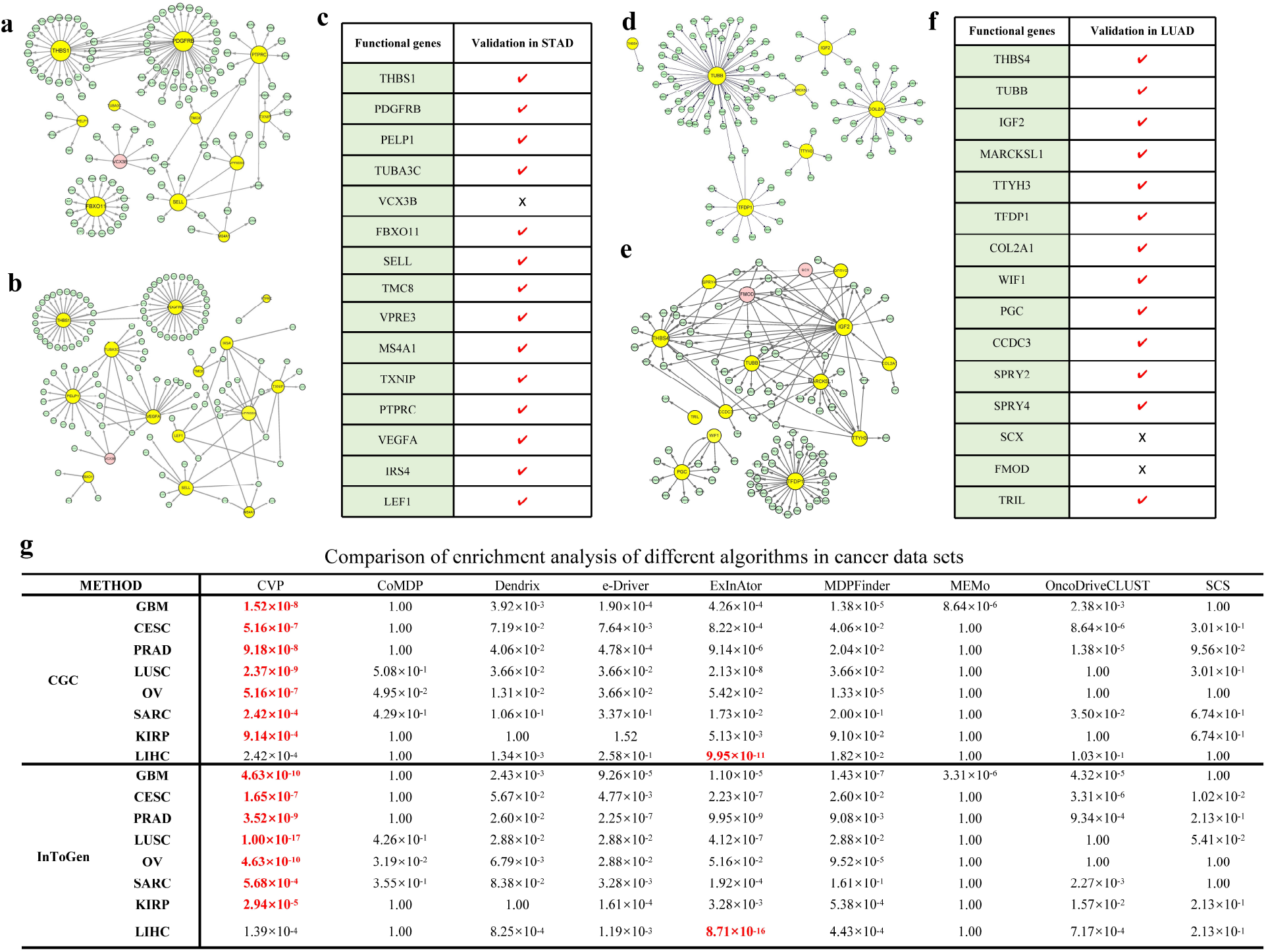
The biological significance of functional genes in cancer networks. a, b are gene regulatory subnetworks for early and late stages of STAD, and the yellow nodes are central genes detected by the CVP algorithm in late cancer stages. c shows the comparison results of 15 functional genes predicted by CVP algorithm in literature verification. d, e are the gene regulatory subnetworks of early and late stages of LUAD, and f is the result of literature verification of 15 functional genes predicted by CVP algorithm. g. The results of enrichment analysis for the predicted driver genes from different algorithms of driver gene inference.

Similarly, based on the CVP algorithm we constructed the regulatory networks for two stages of lung adenocarcinoma (LUAD), the early (Stage I and II) and the late (Stage IV), identifying 15 out-degree hub genes as the functional genes associated with late cancer progression (Fig. 5f and Supplementary Note 9). Based on literature validation, 13 of the 15 central genes we found were strongly associated with the progression of lung cancer (Supplementary Note 9). In lung cancer, the late regulatory network of the 15 functional genes with average out-degree 10 is obviously larger than the early regulatory network with average out-degree 7 (Fig. 5d and 5e). Most of the functional genes gained more downstream regulation genes from early to late in lung cancer (Fig. 5d and 5e). It is consistent with some reported genes, e.g., IGF2 is an important oncogene and its activity was reported to take part in tumor growth and metastasis in lung cancer(*68*) (*69*).

Each type of cancer has its own unique pathogenesis. Using glioblastoma multiforme (GBM) as an example, we investigated the subnetwork of the oncogene tumor protein P53 (TP53) to reveal the specific regulatory pattern of TP53 in GBM. For GBM, we obtained 338 regulatory edges of TP53 in the cancer network, compared to 144 regulatory edges in normal samples (Fig. S14). This revealed the regulatory pattern of TP53 in GBM, indicating that the regulatory molecules of TP53 are significantly increased in cancer. The TP53 may promote tumorigenesis mainly by inducing GBM related edges(*70*), and this is consistent with the reported gain-of-function of TP53 in GBM(*71, 72*).

### Identifying functional driver genes in a network level

The cancer is due to the accumulation of gene mutations, and now it is generally accepted that selective mutations of a small number of genes positively promote the occurrence of cancer, such genes that are called driver genes (*73*). Identifying the genes that drive tumorigenesis and progression in cancer is key to cancer diagnosis and treatment. Despite the rapid development of a large number of algorithms for inferring driver genes, there is still a huge gap in the complete catalogue of driver genes in cancer (*74*).

Here, we identified hub genes from GRNs, which were constructed by the CVP algorithm, perspective to predict driver genes of different cancer. The CVP algorithm is based on strict and consistent baseline analysis, when predicted the drivers. We applied the CVP to eight cancer datasets (GBM, CESC, PRAD, LUSC, OV, SARC, KIRP and LIHC) in TCGA (The Cancer Genome Atlas) to construct the GRNs and all the expression data were normalised using TPM (Transcripts Per Million). We considered that in directed networks, degree centrality alone does not provide a complete picture of the central gene, and often ignores nodes in the network that are critical but have few connected edges. Here, we used the PageRank centrality (*75*), that an algorithm was originally used to rank web page popularity, and the genes with high PageRank centrality were defined as ‘functional driver genes’ involved in the regulation of multiple pathways. Eight GRNs were constructed by the CVP algorithm for the eight cancer datasets, and 100 genes with top PageRank centrality were identified from the GRN as the potential driver genes of each tumor (Tab. S7 and Fig. S13). For validating the importance of these driver genes, we used the potential driver genes to do the enrichment analysis into the CGC(Cancer Gene Census) (*76*) and InToGen databases (*77*). The effectiveness of identifying cancer driver genes is further demonstrated by comparing the results with other 8 algorithms (*78–85*) (Fig. 5g).

Fig. 5g depicts the *p*-values of the nine methods (CVP, CoMDP (*78*), Dendrix (*79*), e-Driver (*80*), ExInAtor (*85*), MDPFinder (*81*), MEMo (*82*), OncoDriveCLUST (*83*) and SCS (*84*)), which were used to predict the driver genes of cancer, for enrichment analysis of the predicted driver genes into CGC and InToGen. The lower *p*-values of the enrichment analysis, reflected the better effect to enrich to the existing cancer genes for the predicted driver genes from different algorithms. Notably, the CVP based method has the best enrichment results in the 7 of 8 cancer datasets (Fig. 5g), with the lowest *p*-values in all methods and orders of magnitude better than the other algorithms. The MEMo is relatively poor at predicting drivers, with a *p*-value of mostly 1, as no intersection of predicted genes and the tumor gene database (CGC or InToGen) or no drivers were predicted due to data limitations (*86*). The fact that the network we have constructed uses only cancer expression data and does not require additional mutation information or methylation data, which means our algorithm is more broadly applicable and is sufficient to explain the importance of these genes to organisms even at the gene expression level. Overall, the network we have built provides a true measure of the complex information in biological systems and identifies potential driver genes and biological functions.

In the LIHC dataset only, our algorithm ranked second (Fig. 5g), which is still a positive result overall. Considering the overall excellent computational results described above, we have reason to believe that our predicted results are strongly likely to contain novel drivers that have not yet been reported. Therefore, we used our inferred driver genes from LIHC dataset to do wet experiments to validate the driver genes and regulatory relationships.

### Functional driver genes inhibited proliferation and colony formation in liver cancer cell

In order to investigate the influence of functional driver genes on tumor growth, we chose the top 10 functional driver genes from LIHC dataset. Then we knocked out the 10 genes (SNRNP200, KHDRBS1, MTPN, MOB1A, XRN2, NCKAP, PPP1R12A, SP1, SRSF10 and RALGAPB) in liver cancer cell line Huh7 using CRISPR-Cas9 (Fig. S15). For the exploration of the proliferative capacity of these cells, colony formation was used to detect the viability of the liver cancer cells after knocking out these genes. Colony formation assay indicated that the formatting of cell colony-forming units was inhibited in KO (knock out) group, providing possible indicators for driver genes detection (Fig. 6a). Compared with control cells, these four factors (SNRNP200, XRN2, RALGAPB and SRSF10) induced about 50% growth inhibition in Huh7 cells (Fig. 6b), so these four gene KO cells were utilized in the following experiments.

**Fig. 6.**
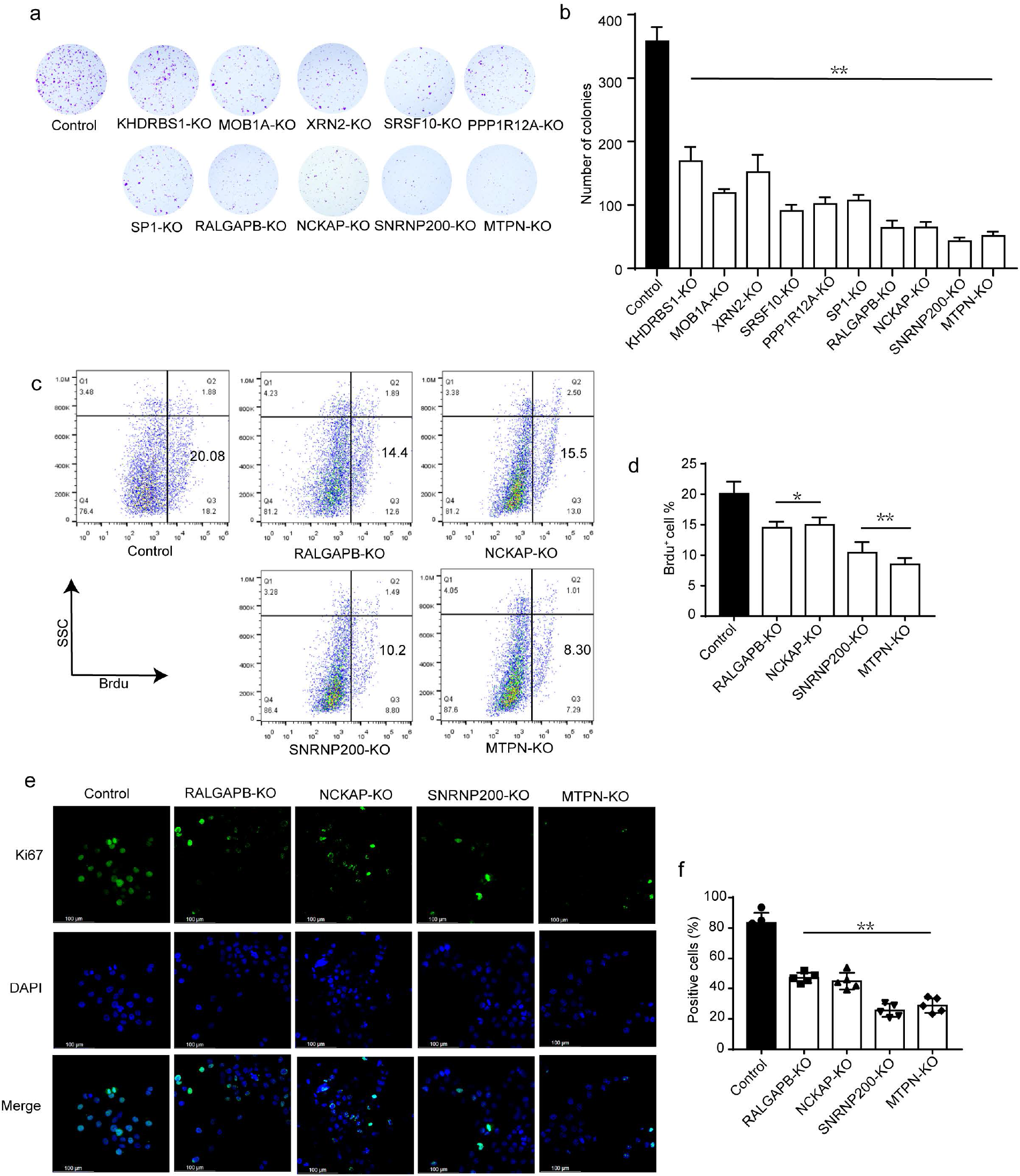
Functional driver genes contributed to proliferation and colony formation in liver cancer cell line Huh7. a. Colony formation of Huh7 cells after knocked out the top ten functional driver genes. b. The statistical analysis of colony formation. c and e were performed in Huh7 cells knocked out SNRNP200, XRN2, RALGAPB and SRSF10 genes. c. Brdu assay of these cells by flow cytometer. d. The statistical analysis of Brdu assay. e. Percentage of Ki67-positive cells by Ki67 staining (×40). Cells were stained with anti-Ki67 antibodies (green) and DAPI to visualize DNA (blue), Ki67 presents (green) and DAPI to visualize DNA (blue), Scale bars: 100 μm. f. The statistical analysis of Ki67 staining. Data are presented as the mean ± SD of three independent experiments. **P < 0.01, *P < 0.05.

Furthermore, Brdu assay was used to measure the proliferation in the above knocked out Huh7 cells. The flow cytometry assay results showed that the Brdu-positive cells were markedly decreased in target-KO group (Fig. 6c). Especially, we found that there are very significant differences in SNRNP200-KO and MTPN-KO cells by statistical analysis (Fig. 6d). In addition, we assessed the immunocytochemistry performance of Ki67, the widely used proliferation marker, to characterize the proliferation activity of tumor cells. The confocal microscopy results indicated that the Ki67 positive percent was reduced significantly in knocked out cells (Fig. 6e), and the results of this statistical analysis for the four groups (RALGAPB-KO, NCKAP-KO, SNRNP200-KO and MTPN, KO) showed that they are a significantly lower than control group (Fig. 6f), which is liver cancer cells line Huh7 cells without any knockout. Taken together these results suggested that the top ten driver gene from LIHC can indeed suppress the proliferation and colony formation in liver cancer cells, and it can validate the power of the driver gene prediction from regulation network by the CVP algorithm.

### The regulation validation in liver cancer cell

In order to validate the causal inference of the CVP algorithm, eight regulations from the regulation network of LIHC were used to verify the accuracy of the CVP algorithm by our biological experiment. To facilitate the biological experiment, we only chose the regulations from the two functional driver genes to their targets in liver cancer. So, four regulations from SNRNP200 to its target genes (MCM2, SMAD2, ORC2 and SMC1A), and other four regulations from RALGAPB to its target genes (REL, MAP2K1, NRAS and MAPK9) were chosen to validate the accuracy of the CVP algorithm by knockout experiment in liver cancer cell line Huh7 cells. When SNRNP200 was knocked out from the liver cancer cell, the gene expression of all the four target genes (MCM2, SMAD2, ORC2 and SMC1A) was strongly downregulated compared with the control cells that is liver cancer cell without SNRNP200 knock out (Fig. S16a). It means that the SNRNP200 can regulate the expression of the four target genes, and it is consistent with the causality from SNRNP200 to the four genes by the CVP algorithm (Fig. S16b).

Meanwhile, after RALGAPB was knocked out from the liver cancer cell, the expression of all four target genes (REL, MAP2K1, NRAS and MAPK9) was downregulated in RALGAPB knock out cells (Fig. S16a). It also means that the RALGAPB can regulate the expression of the four target genes, and it is consistent with the causality from RALGAPB to the four genes by the CVP algorithm (Fig. S16c). As far as we know, the eight regulations are not reported in liver cancer. Then, the eight regulations were predicted by the CVP algorithm in liver cancer and validated by knockout experiment (Fig. S16). It can further reflect the power of the CVP algorithm.

## Discussion

The CVP causality concept with its algorithm was proposed for causal inference on any samples, and its effectiveness was validated by extensive studies of real datasets and also by biological experiments on cancer cell lines. The CVP algorithm quantifies causality between variables based on predictive ability of testing data in a cross-validation manner, which has two major differences from GC, i.e. data type and prediction process. GC is currently well accepted as being effective for inferring causality but on time dependent or time series data, whose prediction of testing data is processed in a time sequential manner. Interestingly, the CVP algorithm also performed better than GC-based method even for the time series data (Tab. S6), which implies that on time series data, our algorithm still has an advantage over the existing algorithms (Supplementary Note 10). A possible reason is the prediction process, i.e. cross-validation based on training and testing groups which are randomly divided, thus ensuring the robust inference of causality. On the other hand, comparing with the algorithms of time-independent data, CVP is able to infer a network with feedback loops common for many real systems, different from SCM mainly for directed acyclic graph without loops.

We show that the CVP algorithm performed better than the existing algorithms by extensive studies on various real cases, including gene regulatory and food chain networks. Our additional biological experiments also proved the predicted “functional driver genes” based on the inferred regulatory network. We highlight three noteworthy aspects. Firstly, a data-driven approach, the CVP algorithm explores the intrinsic characteristics of the data and does not require additional a priori information. Many of the existing algorithms may perform well in some cases but not in other cases, one possible reason is the requirement on some specific hypothesis or some pre-knowledge. Secondly, the CVP algorithm has few parameters to train, keeping consistent parameters under different data backgrounds, thus ensuring the universal operability of the algorithm (Fig. 3 and 4). Thirdly, high false positives have been a thorny problem for causal inference. Our CVP algorithm with condition on other variables (or partial correlation) can eliminate indirect causation by identifying individual causal variables one by one.

The statistical test, e.g., paired *t*-test, can also be used to test the statistical difference of error terms of testing samples for the potential causality. But, statistical test is too time consuming for inferring a large-scale network. In this paper, we used the causal strength *ω_i→j_* to efficiently calculate the difference of error terms of testing samples, which is suitable for a large-scale network.

Causal inference is an importance research topic in various fields including economics, physics, and biology. In biology, identifying causality between molecules benefits deep mechanism study on various diseases or biological processes at a network level. GC test is a classical method to infer the causality between variables, but requires time series data, which limits the application to time independent data. Bayesian network or SCM can infer causality from time independent data, but it generally does not allow the cyclic or loop structure, which is common in the processes of biological regulations. In addition, the computational cost of Bayesian network is generally high, thus unsuitable for analyzing a large-scale biological network. To solve the problems, in this work we developed the CVP causality concept and its efficient algorithm. The CVP causality as well as its algorithm has significant implications for the interpretation of disease mechanisms and biological functions from the perspective of a causal network, and can be applied to analyze a wide range of network problems in various fields.

## Materials and Methods

### The flow of the CVP method

The CVP method quantifies causal relations between variables based on any observed data by cross-validation predictability. Gene expression data with *n* genes and *m* samples is used as an example to explain the implement of the CVP method. Firstly, an initial correlation network is constructed by correlation coefficient (Pearson correlation coefficient or partial correlation coefficient) from the gene expression data, and we then eliminate the edges with low correlation to obtain the correlation network (without directions of edges) from the initial correlation network. Next, the correlation network and the gene expression data are used to infer the causal network (with directions of edges) by the CVP algorithm (Fig. 7a) as the following procedure.

1. Initiation: Each node (or gene) *g_j_* and its first neighbors are considered as a unit (or subnetwork of *g_j_*) in the correlation network (Fig. 7b). We assume that all neighbors of node *g_j_* are the cause of *g_j_*, i.e. the node *g_j_* is assumed to be causally regulated by all its neighbor nodes (Fig. 7b). All of the samples are randomly divided into two groups, i.e. training group and testing group, for cross validation (Fig. 7b).
2. H_1_ model: The node *g_j_* is regressed on all its neighbors in the training samples to obtain regression equation (2) (Fig. 7b), and then the testing samples are used to calculate the error from the regression equation (2) (Fig. 7b). The *e* is the total squared error summation from the regression equation (2) in Fig. 7b for all of the testing samples, i.e. H_1_ model.
3. H_0_ model: For judging the causality of a neighbor node *g_i_* to *g_j_*, the node *g_i_* is removed from the subnetwork of *g_j_* and the gene expression of node *g_i_* is also removed from the subnetwork of node *g_j_* (Fig. 7b). We use the same two groups used in (2) for cross-validation of (1). In other words, for the perturbed subnetwork eliminating *g_i_*, the node *g_j_* is also regressed on all its neighbors except node *g_i_* in the training samples to obtain regression equation (1) (Fig. 7b). Then, the new error is calculated by implementing the regression equation (1) in the testing samples without *g_i_* for predicting *g_j_* (Fig. 7b). The 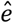 is the total squared error summation from the regression equation (1) in Fig. 7b for all of the testing samples, i.e. H_0_ model.
4. Prediction testing: We used the sum of squared error *e* and 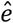 to infer the potential causal regulation from *g_i_* to *g_j_*. If *e* is less than 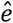, it means that *g_i_* benefits the regression equation (1) to accurately predict the gene expression of *g_j_* in the testing group samples, and *g_j_* can be at least partly explained by *g_i_*, thus the node *g_i_* is the cause of node *g_j_* in this case (Fig. 7b). If *e* is greater than 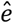, it means that *g_i_* is unsuitable for predicting the gene expression of *g_j_* in the testing group samples, and *g_j_* cannot be explained by *g_i_*, Hence, we consider that the node *g_i_* is not the cause of node *g_j_* in this case (Fig. 7b).

**Fig. 7.**
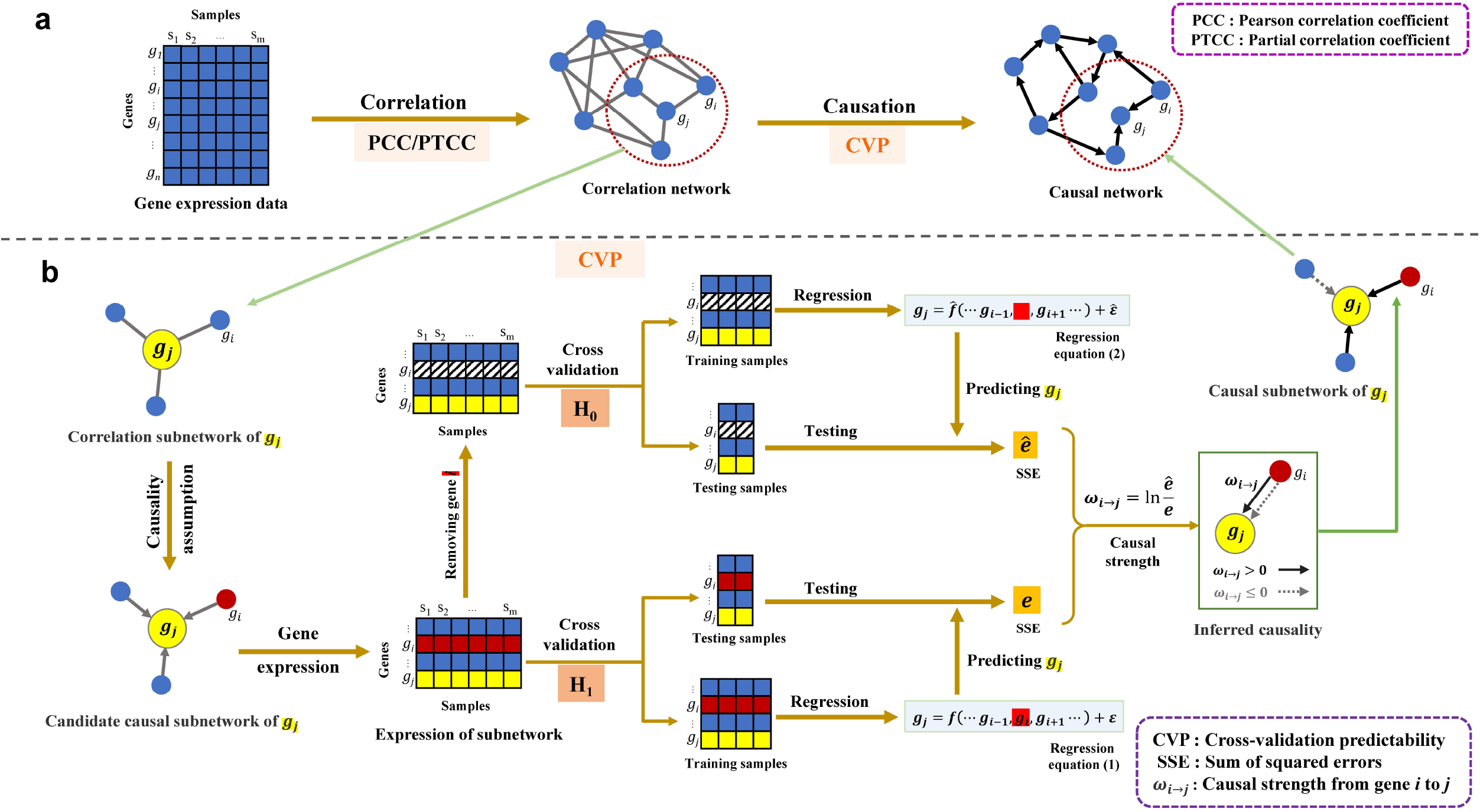
The overall framework of CVP algorithm. a. The general idea of the algorithm, from a correlation network to a causal network. b. The core steps of the CVP algorithm. For the correlation subnetwork of the variable *g_j_*, candidate causal relations are obtained assuming that *g_j_* is modulated by multiple dependent variables. The causal subnetwork of *g_j_* is obtained by cross-validation testing to infer the causal relation of each candidate dependent variable on *g_j_* one by one.

Here, we use causal strength 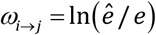 of Eqn. (3) to judge the causality from *g_i_* to *g_j_*, i.e. if 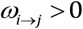, there is causality from *g_i_* to *g_j_*, and if *ω_i→j_* ≤ 0, there is no causality from *g_i_* to *g_j_* (Fig. 7b). After every neighbor node *g_i_* of *g_j_* is evaluated by the corresponding 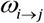 in the subnetwork of *g_j_*, we can obtain the causal subnetwork for *g_j_* (Fig. 7b). Based on the above procedure of CVP algorithm, the causal subnetwork for every node in the correlation network can be inferred one by one, i.e. *j*=1, 2, …, *n*, and then the network combining all *n* causal subnetworks is the inferred causal network with *n* nodes (Fig. 7 and Supplementary Note 1).

### Evaluation indicators

In this study, AUROC (the area under the receiver operating characteristic curve) and AUPR (the area under the precision-recall curve) were used to assess the effectiveness of the algorithm. AUROC is commonly used to evaluate classification models, and its value is the area under the ROC curve. The ROC (receiver operating characteristic) curve is used to weigh the TPR (true positive rate) and FPR (false positive rate) under different decision thresholds, where the horizontal coordinate is the FPR and the vertical coordinate is the TPR. AUROC has a high advantage over unbalanced data with high positive examples, but it does not capture the impact of a large number of negative examples on algorithm performance. Due to the sparsity of GRN, there may be relatively high negative examples, so we consider introducing AUPR. AUPR is the area under the PR (precision-recall) curve, and PR curve trade-offs precision and recall (TPR) at different decision thresholds, with the horizontal coordinate being recall and the vertical coordinate being precision. TPR, FPR, ACC and Precision are calculated as in equations (4).

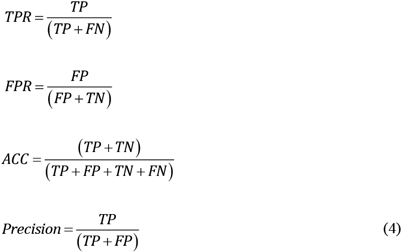

### Data processing and code availability

The data sets covered in this study are available from citations or links in the main text/supporting materials. All cancer datasets are the transcriptome dataset from the TCGA database (https://portal.gdc.cancer.gov/), normalized by TPM (Transcripts Per Kilobase of exon model per Million mapped reads). The codes are available at https://github.com/zyllluck/CVP.

### Knock out

The targeted gRNA oligos were introduced into the lentiviral expression plasmids. The sequences of these oligos are listed in Tab. S7. 4.5×10^6^ HEK293T cells were seeded on a 10 cm dish and transfected with lentiviral expression and packaging plasmids. After another 48 hours, collect medium containing lentivirus particles. The culture medium was refreshed the following morning, and the supernatant containing lentiviral particles was collected after another 48 hours. For experiments, Huh7 cells were transduced with the lentiviral vectors at a multiplicity of infection (MOI) of 0.5 and single colonies were selected with 2 mg/ml puromycin. Gene expression levels in each cell line were examined using Real-Time Quantitative PCR (QPCR).

### Colony formation assay

Gene knocked out Huh7 cell lines were seeded and cultured into the six-well plate at a density of 2 ×10^3^ cells/well in DMEM (Dulbecco’s Modified Eagle Medium) containing 10% FBS (Fetal bovine serum) for 3 weeks to allow colony formation. The plate was washed with cold PBS (Phosphate Buffered Saline). The colonies were fixed by 4% polyformaldehyde at room temperature, washed with water. Next, they were dyed with 1% crystal violet for 30 minutes at room temperature, washed with water several times and dried. Each experiment was done thrice in this study.

### Brdu assay

Brdu staining was used to analyze the cancer cell proliferation according to manufacturer the protocol (Brdu APC flow Kit, 552598, BD). Briefly, knocked out cell lines were incubated with Brdu (10 mmol/L) for 2 hours. The cells were fixed with 4% paraformaldehyde for 20 minutes at room temperature. After permeabilized and staining, cells were collected for flow cytometry analysis.

### Ki67 staining

Huh7 cells were fixed in 1% formaldehyde. Incubate on ice for 5 minutes. Washed twice with PBS containing 0.05% Tween20 and permeabilized with 0.5% Triton X-100 in PBS. Cells stained with the anti-Ki67 antibody (abcam, ab15580, 1:1000) for 1 hour at room temperature. Then the washed cells were stained with Donkey anti-rabbit IgG Alexa Fluor 488 (A32790, Invitrogen, 1: 1000). DNA was stained with DAPI (D9542-1MG sigma).

## Acknowledgements

This work was supported by National Key R&D Program of China (No. 2017YFA0505500), Strategic Priority Research Program of the Chinese Academy of Sciences (No. XDB38040400), NSFC (Nos. 11825102, 12131020, 31930022, 12026608), Zhejiang Provincial Natural Science Foundation of China under Grant No. LZ22C060001, Key Project of Natural Science of Anhui Provincial Education Department (No. KJ2020A0018), Key project of Anhui Finance and Economics University (No. ackyb20015), Project of teaching and research of Department of Education of Anhui Province (No.2020xsxxkc014), Technology Innovation Strategy of Guangdong Province (Nos. 2021B0909050004, 2021B0909060002), Major Key Project of PCL (No.PCL2021A12), and JST Moonshot R&D (No.JPMJMS2021).

