## Supplementary File for "Causal network inference based on cross-validation predictability"

### Supplementary Notes

#### Supplementary Note 1. Implementation process of the CVP algorithm

##### Causality definition and cross-validation predictability method

The causality from CVP method is a statistical concept based on cross-validation prediction for any observed/measured data. Assuming two variables $X$ and $Y$ are observed in *m* samples, we consider that $X$ causes $Y$ if the prediction of the values of $Y$ is improved by including values of $X$ in the sense of cross-validation. Specifically, we assume that a variable set $\{X,Y,Z_1,Z_2,\cdots,Z_{n-2}\}$ includes *n* variables in the *m* samples where $\hat{Z}=\{Z_1,Z_2,\cdots,Z_{n-2}\}$. In other words, $X$ and $Y$ are any two variables among all *n* observed variables. All *m* samples are randomly divided into a training group and a testing group, for the purpose of cross-validation, e.g., *k*-fold cross-validation. To test causal relation from *X* to *Y*, we define a causality framework by constructing two contradictory models $H_0$ (null hypothesis without causality) and $H_1$ (alternative hypothesis with causality) by the same *k*-fold cross-validation and further define its causality strength by the difference between $H_1$ and $H_0$, for quantifying CVP causality.

- $H_0$ model: $$Y=\hat{f}\left(\hat{Z}\right)+\hat{\varepsilon}=\hat{f}\left(Z_1,Z_2,\cdots,Z_{n-2}\right)+\hat{\varepsilon}$$

Train the regression $\hat{f}$ by the training group samples, and then test $\hat{f}$ by the testing group samples in a *k*-fold cross-validation manner. We have the error $\hat{e}_i$ of Eqn. (1) of the main text in the *i*-th cross-validation test by the testing group samples. The total squared testing error is $\hat{e} = \sum_{i=1}^{m} \hat{e}_i^2$ for all *k*-fold cross-validation tests.

- $H_1$ model: $Y = f\left(X, \hat{Z}\right) + \varepsilon = f\left(X, Z_1, Z_2, \cdots, Z_{n-2}\right) + \varepsilon$

  Train the regression $f$ by the training group samples, and then test $f$ by the testing group samples in a *k*-fold cross-validation manner. We have the error $e_i$ of Eqn. (2) of the main text in the *i*-th cross-validation test by the testing group samples. The total squared testing error is $e = \sum_{i=1}^{m} e_i^2$ for all *k*-fold cross-validation tests. If $e < \hat{e}$, then $H_1$ holds, i.e. causal relation from $X$ to $Y$. And if $\hat{e} \le e$, then $H_0$ holds, i.e. no causal relation from $X$ to $Y$.

- Causal strength (CS): $CS_{X \to Y} = \omega_{X \to Y} = \ln \frac{\hat{e}}{e}$

Causal strength is the difference/distance measured between $H_0$ and $H_1$, i.e. $\ln \hat{e} - \ln e$. A paired Student's *t*-test can be used to test the difference between $e$ and $\hat{e}$, i.e. significance test. Here, $f$ is the regression equation for $Y$ fitting $\hat{Z}$ and $X$, and $\hat{f}$ is the regression equation for *Y* fitting $\hat{Z}$ without $X$. In this work, we use the linear regression for both $f$ and $\hat{f}$. The $\varepsilon$ and $\hat{\varepsilon}$ are the error terms of Eqns. (2) and (1) of the main text, respectively. The error is defined as the difference between the predicted value and the true value.

In this paper, we used linear regression to fit the equation from training data. Assuming variables $X\left(x_1, x_2, \cdots x_i, \cdots, x_m\right)$ and $Y\left(y_1, y_2, \cdots y_i, \cdots, y_m\right)$ are observed data in *m* samples, we consider that $X$ causes $Y$ if the prediction of the values of

$Y$ is improved by the inclusion of values of $X$. For exhibiting the inference of observed/measured data, the samples are randomly divided into two groups, training and testing groups for cross-validation.

Seeing that a variable set $\hat{Z}$ { $Z_1\left(z_1^1,z_1^2,\cdots,z_1^m\right)$, $Z_2\left(z_2^1,z_2^2,\cdots,z_2^m\right)$, ... , $Z_{n-2}\left(z_{n-2}^1,z_{n-2}^2,\cdots,z_{n-2}^m\right)$ } includes *n* variables excluding variables $X$ and $Y$ in the *m* samples, and two regression equations were fitted for variable $Y$ on $\hat{Z}$ both with and without $X$ in training group. Here, $\hat{Z}$ presents other elements of correlation or potential causal relation to $Y$ except $X$, and the variable set $\hat{Z}$ can be considered as other factors to affect variable $Y$ except $X$.

For ease of calculation, we used linear regression model to infer the causality from $X$ to $Y$ in this paper, and the linear regression version of the Eqns. (1) and (2) of the main text can be presented:

The linear equation of $\hat{f}$ can be shown as:

$$y_i = \beta_1^{'} z_1^i + \beta_2^{'} z_2^i + \cdots + \beta_{n-2}^{'} z_{n-2}^i + \hat{\varepsilon} = \sum_{j=1}^{n-2} \beta_j^{'} z_j^i + \hat{\varepsilon} \qquad \text{(S1)}$$

And, the linear equation of $f$ can be shown as:

$$y_i = \alpha x_i + \beta_1 z_1^i + \beta_2 z_2^i + \ldots + \beta_{n-2} z_{n-2}^i + \varepsilon = \alpha x_i + \sum_{j=1}^{n-2} \beta_j z_j^i + \varepsilon \qquad \text{(S2)}$$

where $\alpha$, $\beta_j$ and $\beta_j^{'}$ are the coefficients of linear regression, and $1 \le i \le m$. The Eqns. (S1) and (S2) were fitted from the samples in the training group by ordinary least squares to guarantee the dependent variables and the errors to be always orthogonal, and then the samples in the testing group were used to calculated the errors from Eqns. (S1) and (S2), respectively. Hence, linear regression equations (S1) and (S2) are the real model to infer causality in this paper, and they are also a special case of regression equations (1) and (2) of the main text.

The sum of squared errors (SSE) for the testing group are represented as $\hat{e}$ and $e$ from the Eqns. (S1) and (S2) based on the testing samples, respectively. If $H_0$ or $e$ is significantly greater than $\hat{e}$ in the testing group, and it means that Eqn. (S1) is a better regression equation than Eqn. (S2), so Eqn. (S1) is the true regression equation and $X$ does not cause $Y$. If $H_1$ or $e$ is significantly less than $\hat{e}$ in the testing group, it means that Eqn. (S2) is a better regression equation than Eqn. (S1), so Eqn. (S2) is the true regression equation and $X$ causes $Y$.

In this paper, we used a new index $\omega_{i \to j}$ (Eqn. (3)) to represent causal strength and to infer the causality of two variables based on the sum of squares for the errors of regression equations in testing samples (Fig. 7 and Materials and Methods).

**Calculating causal strength**

To measure the causality from node $g_i$ to $g_j$, the errors $\hat{e}_k$ and $e_k$ were obtained for the *k*-th sample based on with and without $g_i$ in the testing samples. In fact, the $\hat{e}_k$ and $e_k$ can form two error vectors including the error of every testing sample based on regression equation (S1) and (S2), respectively, so Student's *t*-test for paired samples can be used to test the significant difference between the two error vectors in testing samples with and without node $g_i$.

In this study, we used a new index $\omega_{i \to j}$ as causal strength to replace the statistical test for inferring the causality from node $g_i$ to $g_j$ (Fig. 7) and the $\omega_{i \to j}$

was defined as equation (S3).

$$\omega_{i \to j} = \ln \frac{\hat{e}}{e} \tag{S3}$$

where $\hat{e}$ is the sum of squared errors of testing samples without node $g_i$ by the regression equation (1) in Fig. 7b, i.e. $\hat{e} = \sum_{k=1}^{m} \hat{e}_k^2$ , and $e$ is the sum of squared errors of testing samples with node $g_i$ by the regression equation (2) in Fig. 7b, i.e. $e = \sum_{k=1}^{m} e_k^2$. In fact, because each testing sample has a pair of $\hat{e}_k$ and $e_k$ , the sums of squared errors $\hat{e}$ and $e$ were used to finally calculate the $\omega_{i \to j}$ after cross-validation.

The *m* is the number of samples, the $\omega_{i \to j}$ was defined as index or causal strength of inferred causality. When the $\omega_{i \to j}$ is greater than a threshold 0, we consider $g_i$ to be the cause of $g_j$ (Fig. 7b), and vice versa.

**Algorithm workflow**

The algorithm consists of two processes: (i) The correlation between two variables was calculated by Pearson correlation coefficient (*PCC*) and partial correlation coefficient (*PTCC*), to eliminate the indirect influence of genes and calculate the direct correlation among genes. (ii) Cross-validation predictability (CVP), to screen out the optimal set of regulation for each target gene (Fig. 7 and Materials and Methods).

The CVP algorithm calculates the direct causality between genes, and constructs the optimal regulatory network. The algorithm is mainly divided into three steps. Firstly, separate the subnetwork of every gene with first-order neighbors from the correlation network. Secondly, calculate the prediction errors for regression equation eliminating each neighboring gene by cross-validation. Finally, compare the prediction errors of different regression equation to obtain the optimal causal sub-network for every gene (see Fig. 7 and Materials and Methods).

**Pearson correlation coefficient**

The Pearson correlation coefficient (*PCC*) is used to measure the strength of correlation between two variables (https://en.wikipedia.org/wiki/Pearson_correlation_coefficient) and the *PCC* was calculated using the extension module 'Pandas' (https://pandas.pydata.org/) of Python programming language. We constructed the correlation network using *PCC*, with network nodes denoting genes, and the edges represent the strength of the correlation between genes. Obviously, the value of correlation is influenced by many factors, such as sample size, sequencing depth, etc. In order to reduce the impact of technical factors on different data sets, we used adaptive rather than fixed thresholds. If the feature dimension is lower than 1000, the correlation network chooses to retain all the edges, otherwise, the highly correlated edges with the top 100 average degree will be retained.

The purpose of *PCC* for building the correlation network is to reduce the computational cost before using *PTCC* to obtain the direct correlation network.

**Partial correlation coefficient**

In statistics, partial correlation coefficient (*PTCC*) is used to evaluate the directly correlation between two variables, which is the correlation network obtained after excluding a set of random variables (https://en.wikipedia.org/wiki/Partial_correlation#Using_linear_regression). In this paper, the 'Pingouin' extension package (https://pingouin-stats.org/) of Python programming language is used to calculate the *PTCC* among genes. The partial correlation network was constituted by the edges with the *p*-value of the *PTCC* less than 0.1. The principle of *PTCC* is followed as:

Given a variable set $Z\{Z_1, Z_2, \cdots, Z_{n-2}\}$, the partial correlation coefficient between two variables/genes $X$ and $Y$ controlling $Z$ can be represented as $PTCC(XY/Z)$. In order to calculate the *PTCC* between gene $X$ and $Y$, the $X$ and $Y$ were separately regressed on variable set $Z$, and the regression equations were shown below, respectively.

$$X = \sum_{l=1}^{n-2} k_{1l} Z_l + b_1 + e_X \tag{S4}$$

$$Y = \sum_{l=1}^{n-2} k_{2l} Z_l + b_2 + e_Y \tag{S5}$$

where $k_{1l}$ and $k_{2l}$ are regression coefficients in Eqns. (S4) and (S5), $e_X$ and $e_Y$ are error terms in Eqns. (S4) and (S5), and the value of $l$ is between 1 and $n$-2. In this case, the $PTCC(XY/Z)$ is the value of the Pearson correlation coefficient between the errors $e_X$ and $e_Y$ as $PCC(e_X, e_Y)$.

The control variables set $Z$ contains the genes that link to both $X$ and $Y$ at the same time on the correlation network of *PCC*.

**Cross-validation predictability**

The cross-validation predictability (CVP) is the core concept of our algorithm. The CVP algorithm inputs target gene $g_j$ and all potential regulators and outputs $g_j$ and its inferred regulators (Fig. 7b). Firstly, all potential regulators were arranged in ascending order of correlation coefficient, and the regulators were obtained as $G\{g_1,g_2,\ldots,g_i,\ldots,g_n\}$. When executing the algorithm, regulators were screened in order of *PTCC* from low to high, and the initial screening gene was $g_1$, that is, the least important gene for $g_j$. For any regulator $g_i$, whether to delete or retain the edge from $g_i$ to $g_j$ was judged according to $\omega_{i\to j}$. When $\omega_{i\to j}>0$, the control relationship was retained, otherwise, the edge was removed. Each iteration of the algorithm is based on the network structure of the previous step. After all the regulators were screened, the optimal regulatory subnet of target gene $g_j$ was generated (Fig. 7b).

#### Supplementary Note 2. Algorithm parameters

To validate CVP algorithm, five popular methods for constructing regulation network were used to compare the performance of CVP, including Nonlinear ODEs [1], GENIMS [2], PLSNET [3], GENIE3 [4], TIGRESS [5]. The default parameters were used for the most software packages of the five methods. For example, the integration parameters of PLSNET, GENIE3, GENIMS were set to 1,000. GENIE3 analysis was based on random forest feature selection, where four algorithms were used by considering four parameters: RF, ET, all, and sqrt. The PLSNET method differs from the others, and a parameter K is more difficult to determine with different network sizes. The method for selecting the regulator followed that of GENIE3, where the main parameters of PLSNET were set to *nfac*=3 and $K = \sqrt{p}$ . PLSNET was ran in MATLAB, while GENIMS and TIGRESS were used the R package provided in the original methods. GENIE3 and Nonlinear ODEs were run with python code.

#### Supplementary Note 3. Inferring causality of three nodes from simulated data

In most real cases, it is not possible to detect true cause and effect due to objective limitations [6]. Therefore, inferring causality from the statistical properties of observed data is crucial, and a very challenging task [7]. Here, we used three nodes to construct five causality cases to simulate different causality among variables (Fig. S2), and five simulation datasets were generated from the five causality cases (Tab. S1). Then, the CVP algorithm was used to infer the causality from the five simulation datasets.

The simulation data of linear causality for two variables can be generated by the below equation:

$$Y = aX + \varepsilon \tag{S6}$$

Where, $Y$ is generated from both $X$ and $\varepsilon$, and $X$ and $\varepsilon$ were randomly generated and orthogonality ( $\varepsilon \perp X$ ). Strictly speaking, only one of the variables $X$ and $\varepsilon$ is from Gaussian distribution at most. So, the regression noise should be independent of the predictive variables, and then it can lead to the asymmetric causality between $X$ and $Y$ [7].

In order to obtain the discernable causal connection, the simulation data were generated based on above principle for all 5 structures or cases of the variables $X,Y,Z$ (Fig. S2a-S2e and Tab. S1), and the independent variable(s) sampled from the Gaussian distribution and the noise from the uniform distribution (Tab. S1). We randomly generated 1000 samples for each structure or case in Fig. S2a-S2e, and the sample size is 1000 to ensure the robustness of the sampling results. The Fig. S2f-S2j shows the inference results of the 5 structures or cases from the simulation data by the CVP algorithm, and it is worth noting that the CVP algorithm can effectively identify the true causality and eliminate the false causality (Fig. S2f-S2j). Obviously, for the non-existent causality, the CVP algorithm can obtain the steady value of 0 (Fig. S2f-S2g). And we emphasize that our algorithm can effectively distinguish the real causality structure, whatever it is indirect causation caused by cascade or false causation caused by confounding factors (Fan-out) or collision factors (Fan-in) (Fig. S2).

Specifically, the CVP algorithm is able to accurately determine the true causality and eliminate spurious relationships. And the CVP algorithm can also effectively eliminate indirect causality, such as the spurious connections generated due to cascading (Fig. S2c). In addition, by simulating the causality of different structures, the CVP algorithm is feasible and robust in the basic causality structure, providing sufficient theoretical support for the subsequent analysis of reconstruction of large complex network (Fig. S2).

#### Supplementary Note 4. Identifying causality of multivariate from simulated data

The CVP algorithm assumes that any one variable is regulated by multiple variables in a system. To verify the rationality of hypothesis, 9 structures or cases of causality (Fig. S3a) were used to separately generate simulation data (Tab. S2), and each simulation dataset with 1000 samples was randomly generated for each structure or case (Fig. S3a and Tab. S2). Then the causalities of the 9 structures or cases were inferred by the CVP algorithm from the simulation data (Fig. S3b-S3c). The comparisons between the real causality and the inferred causality (Fig. S34b-S3c) can reflect that the power of the CVP algorithm can accurately recover the real causality from simulation data.

To test the robustness of the CVP algorithm, we constructed a causality structure including a center node and 10 neighbor nodes, and then deleted edges one by one and estimated the sensitivity of the CVP algorithm on the causality structures. Fig. S3b shows the real and inferred causalities in the 9 cases from the CVP algorithm, and the horizontal axis is the real and inferred causality in different cases. The T1…T9 represent the real causality of case1…case9, and the P1…P9 represent the inferred causality of case1…case9. Obviously, CVP can accurately identify the real causal edge with a probability of 100% (Fig. S3c). For the deleted edges in different structures or cases (Fig. S3a), that is non-existent edges, the probability of accurately inference using the CVP algorithm is also close to 100% (Fig. S3b-c). We listed all the real structures, and the average accuracy computed by the CVP algorithm on each structure or case repeated 1000 times and was shown in the Fig. S3c. Therefore, our algorithm is significantly effective in the multi-to-one structural assumptions, and can find the potential "cause" variables for any "effect" variable, thus CVP algorithm is expected to be widely applied in other complex data.

#### Supplementary Note 5. Performance of more simulation data from DREAM challenges

The DREAM challenges (https://dreamchallenges.org) include golden standard regulation networks that were experimentally obtained in yeast or E. coli, and corresponding simulation dataset which can be used to validate the performance of algorithms. In addition, both non-time series/steady-state data (e.g., Network1 and Netwotk3 in Fig. 2) and time-series data (e.g., Network2 and Network4 in Fig. 2) are included, allowing for the collection of corresponding data for different algorithms. DREAM3 and DREAM4 are sub-projects of the DREAM challenges, and we selected two data of size 10 from the two sub-projects respectively for the verification of the CVP algorithm. In order to ensure the universality of the CVP algorithm, we selected non-time series/steady state data and time series data in the two subprojects. Network1, that belongs to DREAM3 challenge and comes from the steady-state (non-time series) level of wild-type of the yeast and heterozygous knock-down strains of each gene, included 10 nodes and 22 regulation edges (Fig. 2a and Tab. S3).

Network2 is time series data from DREAM3 containing the time course for the network recovered from an external perturbation of yeast, consisting of 10 nodes with 25 edges (Fig. 2b and Tab. S3). Network3 (Fig. 2c) and Network4 (Fig. 2d) are networks from DREAM4, where Network3 comes from a non-time series data and contains 10 nodes with 16 edges, and Network4 comes from a time series data and consists of 10 nodes with 13 edges (Tab. S3).

In order to compare the sensitivity and accuracy of the algorithm at different scale data, we also quantitatively evaluated the performance of other algorithms (CVP, Nonlinear ODEs [1], GENIMS [2], PLSNET [3], GENIE3 [4] and TIGRESS [5]) on networks with 10, 50 and 100 nodes, in addition to the simulation data included in the main text. We benchmarked using DREAM Challenges [8,9] data and compared ROC and PR curves of different algorithms for a 10-nodes dataset. For large size networks (50 and 100 nodes), we focused on evaluating the inferred network structure, and used accuracy to evaluate the effectiveness of these algorithms.

In the comparison of the six extended datasets (Tab. S3) of 10 nodes, the CVP algorithm always consistently outperformed the other algorithms in both the ROC curve and the PR curve, with the highest AUROC values and PR values (Fig. S5). The performances of the other algorithms were not stable, for example, the TIGRESS algorithm ranked second in the fourth dataset (Fig. S5d and S5k) in terms of AUROC and AUPR values, but was not as good in the other datasets. At the same time, a high AUROC value for the algorithm is not necessary with a good AUPR value. For example, in the fifth dataset (Fig. S5e and S5l), GENIE3_ET_all performed similarly to CVP and tied for first place in the ROC curve with AUROC 0.72 (Fig. S5e), but the AUPR value 0.28 was much lower than 0.34 of CVP (Fig. S5l). Therefore, the CVP algorithm is robust and effective in 10 nodes.

We inferred causal networks for 50 and 100 nodes by simulation data to compare with real networks. Fig. S6 and S7 respectively reflected the accuracy of inferred networks for 50 nodes and 100 nodes. On the whole, even the same algorithm has very different results on different networks[10]. However, the CVP has the highest accuracy in all 30 data sets, with the accuracy of more than 50% data to be above 0.9 and the average accuracy to be 0.89. For some networks that are not easy to explain, such as Net 9, 10, 13, 15, 25 and 30 (Tab. S3), although the accuracy of all the algorithms is low, the CVP algorithm is at least 10% more accurate than the second ranked algorithm (Fig. S6 and Fig. S7). Especially for Net 25 (Tab. S3) where the accuracy of the other five algorithms is around 0.57, but the CVP algorithm has a more prominent advantage with an accuracy of 0.91 (Fig. S7).

#### Supplementary Note 6. Comparison of the additional performance in the IRMA data

In addition to the assessment results in the main text, we have still conducted comparative analyses of other indicators for the IRMA dataset [11], as detailed in Tab. S4. We also considered ACC (accuracy), AUROC, AUPR, TP (Ture positive), FP (False positive), TN (true negative) and FN (false negative), the seven indexes to evaluate the nine algorithms.

The IRMA network is a synthetic network embedded in Saccharomyces cerevisiae, cultured in vivo to obtain ground biological data, and is widely used as benchmark data in inferential modelling. This network contains five genes and eight regulatory edges, whose structure is known (Fig. 2i). The IRMA network is divided into two interference experiments: switch on and switch off. In the switch on experiment, the cells switch from glucose to galactose. In the switch off experiment, the experiment is reversed. We chose the third switch on experiment, which contains five genes and eight regulatory edges for a time series data with 9 time points, to carry out algorithmic reasoning.

The inference results show that our CVP algorithm presents an overall superiority. Regardless of the choice of any index, the CVP results were the best, and can identify the closest network to the true network (Fig. 2i-l and Tab. S4). Specifically, CVP inferred 6 edges with only 1 false positive edge. Similarly, the TIGRESS algorithm also performed well, and it also inferred 6 edges but with 2 false positives. Other methods had even less satisfactory results, with GENIMS inferring 7 edges and 5 false positives (Tab. S4).

#### Supplementary Note 7. Comparison of the additional performance in the SOS data

The SOS DNA repair network from experiment in *E. coli* [12,13] is the most commonly used as a benchmark for real datasets (for a network with 9 genes and 24 edges), which is also a relatively complex nonlinear network at present (Fig. 3i). The results are significantly difference from those of the IRMA data (Fig. S8), with the SOS data showing significantly lower values both AUROC (Fig. 3a and Fig. S9a) and AUPR (Fig. 3e and Fig. S9b), which is inextricably linked to the complexity and sparsity of the real network and reflects the difficulties of real network inference. However, for our algorithm, the evaluation index did not differ too much with the results of IRMA data, and the CVP algorithm identified 16 edges and just three false positives, a rare result for inferred networks on this data [14-16] (Fig. 3j and Tab. S5). Meanwhile, the accuracy of CVP is 0.81 (Tab. S5), far higher than other algorithms (Tab. S5), and the inferred causal network is the closest to the real regulatory network (Fig. 3i-3l). As for the other algorithms, the false positive edges are higher than the real positive edges, which reflects the complexity of the network and the difficulty of reconstruction task (Tab. S5).

#### Supplementary Note 8. Comparison of the additional data in the TCGA database

In the main text, we only compared the performance of eight different algorithms on the BLCA dataset, and we also collated results from other four tumor datasets, such as Uterine Corpus Endometrial Carcinoma (UCEC), Liver hepatocellular carcinoma (LIHC), Pancreatic adenocarcinoma (PAAD), and Stomach adenocarcinoma (STAD). All data were collected from the Cancer Genome Atlas (TCGA, https://www.cancer.gov/about-nci/organization/ccg/research/structural-genomics/tcga) database with level 3 data set, normalized by transcripts per million (TPM) expression data and ground truth regulatory networks were obtained from the corresponding tumor pathways on the KEGG database, where UCEC, LIHC, PAAD and STAD correspond to Endometrial cancer pathway (hsa05213), Hepatocellular carcinoma pathway (hsa05225), Pancreatic cancer pathway (hsa05212), and Gastric cancer pathway (hsa05226), respectively. Overall, the AUROC values of the nine algorithms for the four datasets (Fig. S11) were slightly lower than the AUROC value of BLCA (Fig. 3d), but the performance of the CVP algorithm was still better than other algorithms (Fig. S11). The main reason for the low AUROC values may be that the KEGG pathway only contains a part of the regulatory relationship, and there are many potential regulatory edges that have not yet been identified.

#### Supplementary Note 9. The identified key genes in the late gastric and lung cancers

The CVP algorithm was used to infer the regulatory network of the late gastric cancer, and the top 15 hub genes were identified in the regulation of the late gastric cancer. Notably, it has been demonstrated through literature validation that 14 of the 15 hub genes have been used as association genes for gastric cancer and play a crucial role in gastric cancer[17], namely THBS1[18], PDGFRB[19], PELP1[20], TUBA3C[21], FBXO11[22], SELL[23], TMC8[24], VPRE3[25], MS4A1[26], TXNIP[27], PTPRC[28] VEGFA[29], IRS4[30] and LEF1[31](Fig. 5c). We have described the mechanism of action of certain hub genes in gastric cancer, such as THBS1 (In the latest studies, the methylation state of THBS1 in gastric cancer and its clinical pathological significance were discussed, and THBS1 was proven to be a new marker that predicts gastric cancer[32]. At the same time, the existing studies have shown that THBS1 is positively correlated with high expression in gastric cancer, and is associated with anti-cancer drugs, such as Orshaliplatin [17]), PDGFRB (The expression level of PDGFRB in gastric cancer is closely related to the overall survival rate and can be followed as a close indicator of survival in gastric cancer patients, and this gene can be used as a potential marker for gastric cancer [33]), PELP1(PELP1 encodes a transcription factor whose co-activation by other hormone receptors has been shown to be directly linked to the oncogenic process in a number of cancers, particularly hormone-dependent cancers. In gastric cancer, the expression of this gene is positively correlated with tumor tissue grade and TNM stage, and is an oncogene in gastric cancer. PELP1 also controls the activation of the c-Src-PI3K-ERK pathway, which is the main possible cause of carcinogenesis and may be a potential therapeutic target for chlorpromazine in gastric cancer[20] and TUBA3C (In pan-cancers such as gastric and oesophageal cancers, TUBA3C is mutated in a flanking cationic residue within the protein kinase PKC common site. This disorder may result in partial or complete impairment of PKC phosphorylation, a condition that promotes proliferative tumors [21]. VCX3B is part of the VCX family, a gene with multiple members on both the X and Y chromosomes and is a potential biomarker for gastric cancer that we have discovered. DNA methylation is widely considered to have great potential as a biomarker for the treatment of cancer[34]. However, the function of VCX3B is unclear and has not been used in clinical studies. Comparing the VCX3B homologous sequence of the VCX family, we suggest that the increase of VCX3B may improve the stability of certain mRNAs and control their decapitation [35], which may provide an effective strategy for the targeted treatment of visceral sarcoma by studying its associated functional regulation in visceral sarcoma. Therefore, CVP algorithm can not only accurately identify biomarkers of gastric cancer, but also effectively identify the directed network information, providing key information for early intervention and monitoring treatment of gastric cancer in the further research.

The CVP algorithm constructs a directed cancer network and we identify 15 pivotal genes in advanced lung adenocarcinoma data (Fig. 5f), 13 of which have been shown to be associated with lung cancer, such as THBS4[36] , TUBB[37], IGF2[38], MARCKSL1[39], TTYH3[40], TFDP1[41], COL2A1[42], WIF1[43], PGC[44], CCDC3[45], SPRY2[46], SPRY4[47] and TRIL[48]. Only the regulatory mechanism of SCX and FMOD in lung adenocarcinoma is not clear. It shows that our study is clinically important to accurately identify biomarkers of lung adenocarcinoma and may provide new targets for the treatment of lung adenocarcinoma. Among these, MARCKSL1, a family member of MARCKS, is closely associated with poor prognosis in lung adenocarcinoma[39]. Metastasis is currently the leading cause of cancer death, and silencing MARCKSL1 inhibits metastasis and invasion of lung adenocarcinoma cells and can be used as a biological target in combination with other drugs to treat lung adenocarcinoma[49]. Meanwhile, the genes SCX and FMOD we discovered are considered to be potential functional genes related to lung adenocarcinoma. However, the mechanism of action of these two genes in lung adenocarcinoma is not clear at present, which needs to be verified by biological experiments in the later stage. Based on the overall performance of the CVP algorithm, we have reason to believe that these two genes may be potential functional genes in lung adenocarcinoma.

#### Supplementary Note 10. Causal effect estimation for time series data

We proposed the CVP algorithm, a model-free method based only on observation data to identify causality. The mechanism of the CVP algorithm to infer causality between variables is based on the predictability for observation data. In order to verify whether the CVP algorithm can still work well based on the time series data, we compared the CVP algorithm with the time-series based methods, *e.g.*, Granger causality (GC) [50-52] and partial correlation combined with GC(PTCC+GC). We found that the results for either GC or PTCC+GC were not robust across the three time series datasets (DDGni, HeLa and IRMA), with GC achieving an AUROC value of 0.63 in the DDGni dataset and just under 0.5 in the other two datasets (Tab. S6). The PTCC+GC achieved an AUROC value of 0.65 in the IRMA dataset, while the performance in the HeLa data was also unsatisfactory (Tab. S6). The AUROC value of the CVP's result is around 0.8 for the three time series datasets (Tab. S6), and the CVP method is better than both GC and PTCC+GC based on the time series datasets. Inferring causality from time series data considers mainly the chronological order to determine the direction of the causality, but it is difficult to infer causality for non-time series data due to the lack of time sequence or order information. However, the CVP algorithm incorporates the idea of cross-validation to use a prediction to decide the direction of an edges or a causality in a regulation network, then it fully samples the data and ensures that the prediction error is reduced to a minimum. As a result, our algorithm is more robust and the resulting network is close to the true causality.

#### Supplementary Note 11. Data availability

The data of BLCA, UCEC, PAAD, LIHC and STAD were obtained in TCGA database(https://www.cancer.gov/about-nci/organization/ccg/research/structural-genomics/tcga), but the database did not provide the truth network structure. So, we used the pathways of corresponding cancers from KEGG database (https://www.genome.jp/kegg/pathway.html) regarding as the ground truth of the five datasets.

The data on the number of new coronavirus infections in Japan by region downloaded from https://toyokeizai.net/sp/visual/tko/covid19/.

### Supplementary Figures and tables Legends

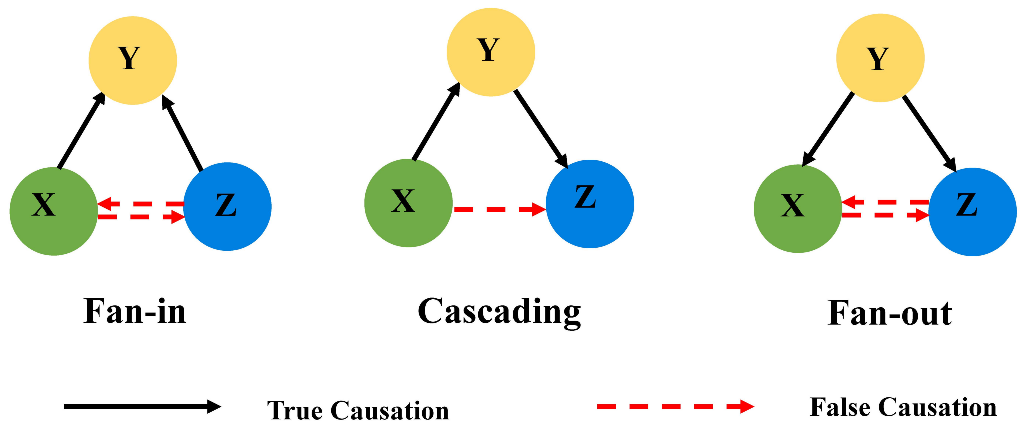


**Fig. S1. Three basic causal networks for node Y.** The solid black lines representing true causality and dashed red lines representing mistakable edges that are susceptible to error detection. Node Y is considered as the center of the three causal networks. Fan-in indicates that X and Z are

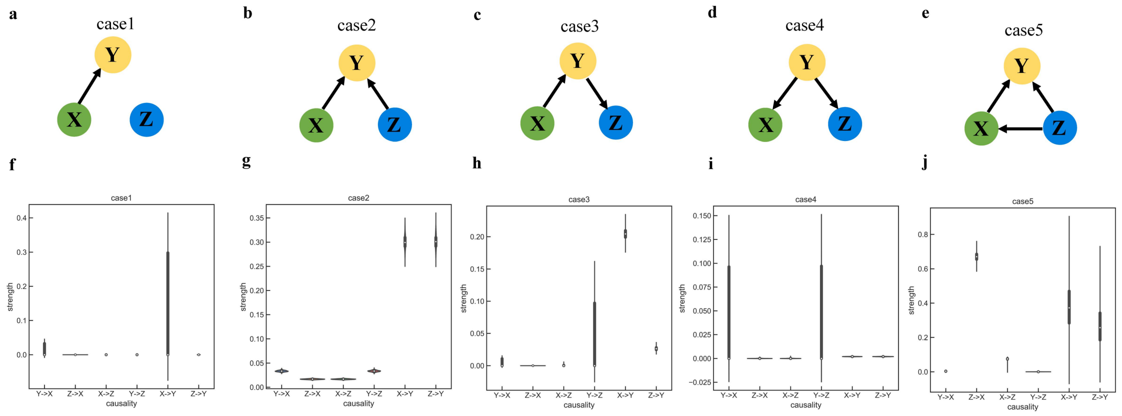


common causes of Y. Fan-out means Y is the common cause of X and Z.

**Fig. S2. Detecting causal structures of three variables.** The graphical model of the causal hypothesis for the three variables aims to estimate the causal effects of variables $X$ and $Z$ on $Y$ from the observed data. a-e, five typical causal structures. f-j, show the inferred causality corresponding to a-e respectively. The horizontal coordinates show the causality of the two variables and the vertical coordinates show the causal strength.

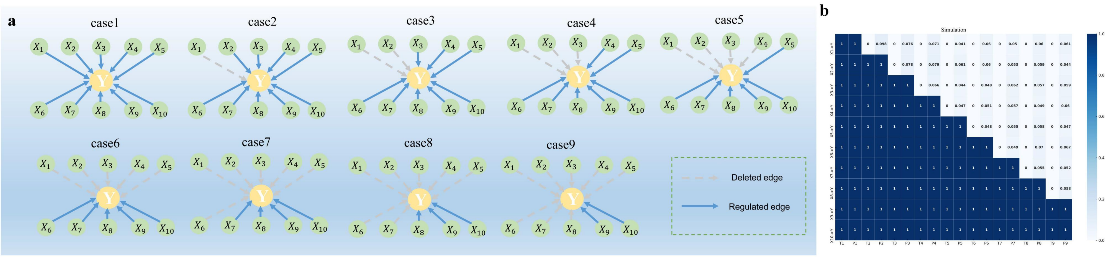


c

The average accuracy from the CVP algorithm under 1000 simulated data in nine cases

| cases | 1 | | 2 | | 3 | | 4 | | 5 | | 6 | | 7 | | 8 | | 9 | |
|---|---|---|---|---|---|---|---|---|---|---|---|---|---|---|---|---|---|---|
| | T1 | P1 | T2 | P2 | T3 | P3 | T4 | P4 | T5 | P5 | T6 | P6 | T7 | P7 | T8 | P8 | T9 | P9 |
| $X_1 \to Y$ | √ | 1.0 | × | 0.902 | × | 0.924 | × | 0.929 | × | 0.959 | × | 0.94 | × | 0.95 | × | 0.94 | × | 0.939 |
| $X_2 \to Y$ | √ | 1.0 | √ | 1.0 | × | 0.922 | × | 0.921 | × | 0.939 | × | 0.94 | × | 0.947 | × | 0.941 | × | 0.956 |
| $X_3 \to Y$ | √ | 1.0 | √ | 1.0 | √ | 1.0 | × | 0.934 | × | 0.956 | × | 0.952 | × | 0.938 | × | 0.943 | × | 0.941 |
| $X_4 \to Y$ | √ | 1.0 | √ | 1.0 | √ | 1.0 | √ | 1.0 | × | 0.953 | × | 0.949 | × | 0.943 | × | 0.951 | × | 0.94 |
| $X_5 \to Y$ | √ | 1.0 | √ | 1.0 | √ | 1.0 | √ | 1.0 | √ | 1.0 | × | 0.952 | × | 0.945 | × | 0.942 | × | 0.953 |
| $X_6 \to Y$ | √ | 1.0 | √ | 1.0 | √ | 1.0 | √ | 1.0 | √ | 1.0 | √ | 1.0 | × | 0.951 | × | 0.930 | × | 0.933 |
| $X_7 \to Y$ | √ | 1.0 | √ | 1.0 | √ | 1.0 | √ | 1.0 | √ | 1.0 | √ | 1.0 | √ | 1.0 | × | 0.945 | × | 0.948 |
| $X_8 \to Y$ | √ | 1.0 | √ | 1.0 | √ | 1.0 | √ | 1.0 | √ | 1.0 | √ | 1.0 | √ | 1.0 | √ | 1.0 | × | 0.942 |
| $X_9 \to Y$ | √ | 1.0 | √ | 1.0 | √ | 1.0 | √ | 1.0 | √ | 1.0 | √ | 1.0 | √ | 1.0 | √ | 1.0 | √ | 1.0 |
| $X_{10} \to Y$ | √ | 1.0 | √ | 1.0 | √ | 1.0 | √ | 1.0 | √ | 1.0 | √ | 1.0 | √ | 1.0 | √ | 1.0 | √ | 1.0 |

**Fig. S3. Modelling the different causal structures of ten variables.** a, nine multivariable causal structures. b, the heat map of comparing real causality and CVP algorithm inference. The vertical axis corresponds to the causal connection structure between nodes in Fig. S3a, and the horizontal axis represents the real connection on T1(True case1), T2(True case2), T3(True case3), T4(True case4), T5(True case5), T6(True case6), T7(True case7), T8(True case8), and T9(True case9), and the inferred accurate probability of causal structures on P1(Inferred case1), P2(Inferred case2), P3(Inferred case3), P4(Inferred case4), P5(Inferred case5), P6(Inferred case6), P7(Inferred case7), P8(Inferred case8) and P9(Inferred case9) for the nine cases. The closer the colors of the real structure and inferred results are, the higher the accuracy is. c, the results of inferring the causality under the nine cases. A check mark indicates an existing edge and a fork indicates no edge in T1…T9. And the numerical values indicate the average accuracy from the CVP algorithm under 1000 simulated data in P1…P9.

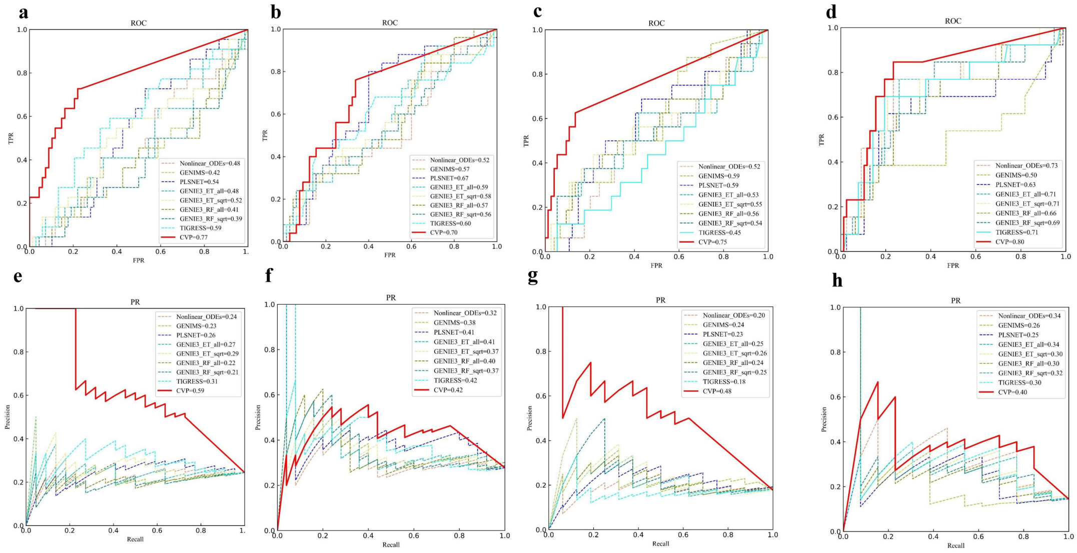


**Fig. S4. Four networks from the DREAM Challenges**. a,b,c,d is the ROC curves for Network 1, Network 2, Network 3 and Network 4 in the main text, and e,f,g,h are the PR curves for Network 1 Network 2 Network 3 and Network 4.

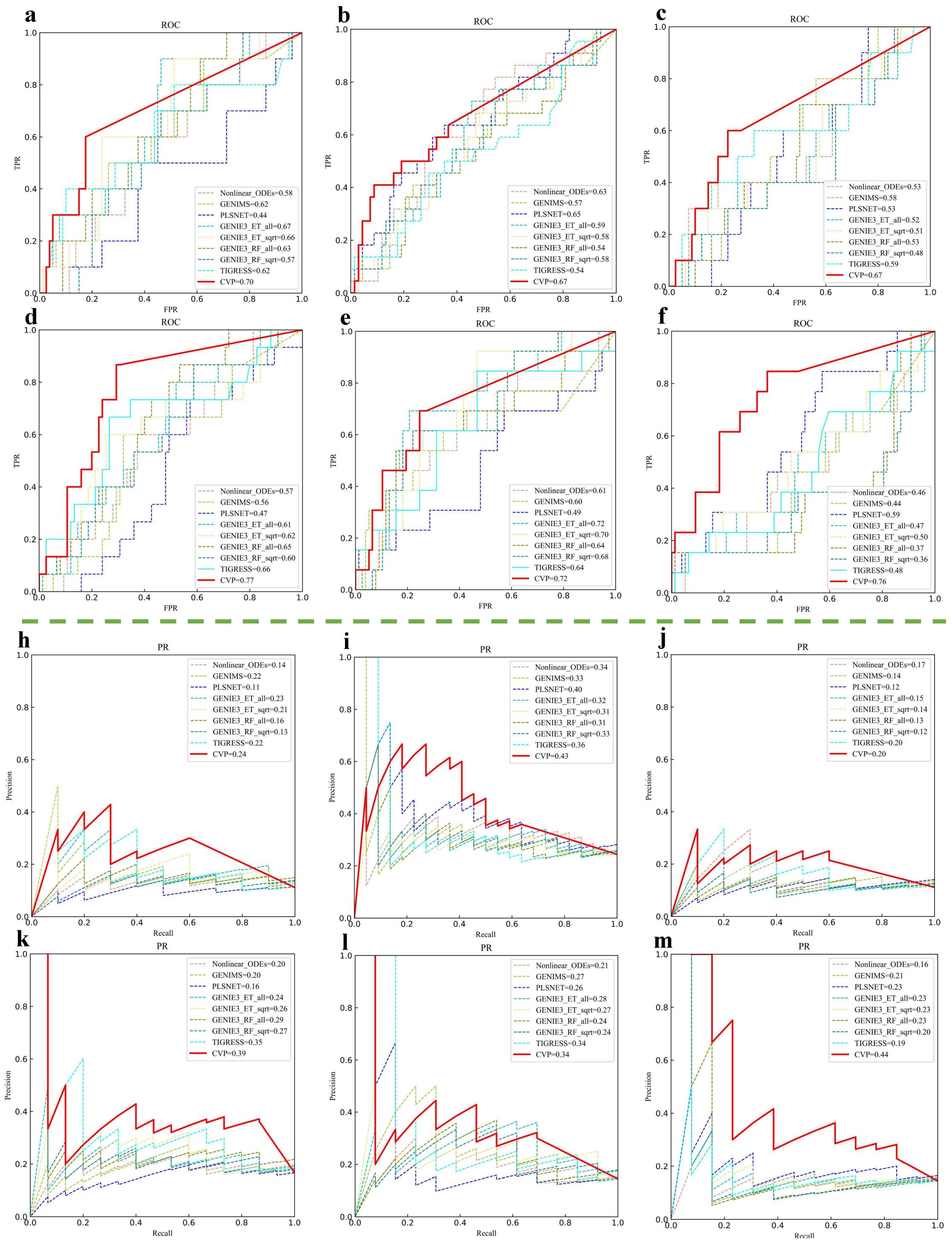


**Fig. S5. Additional comparison outside the main text on ten nodes from DREAM challenges.**

In the six datasets of size 10, a, b, c, d, e and f correspond to the ROC curves of the 9 algorithms and g, h, i, j, k and l correspond to the PR curves of the 9 algorithms.

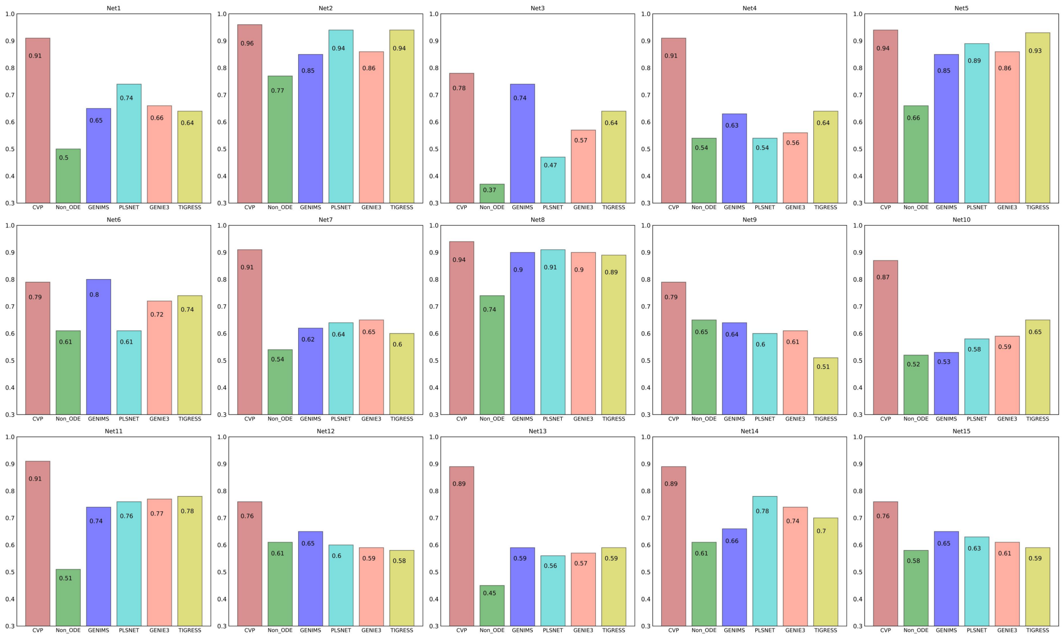


**Fig. S6. Comparison of the CVP algorithm with other eight algorithms for benchmark data of size 50 nodes.** The horizontal coordinate of each subplot represents the nine algorithms and the vertical axis corresponds to the accuracy rate of each network.

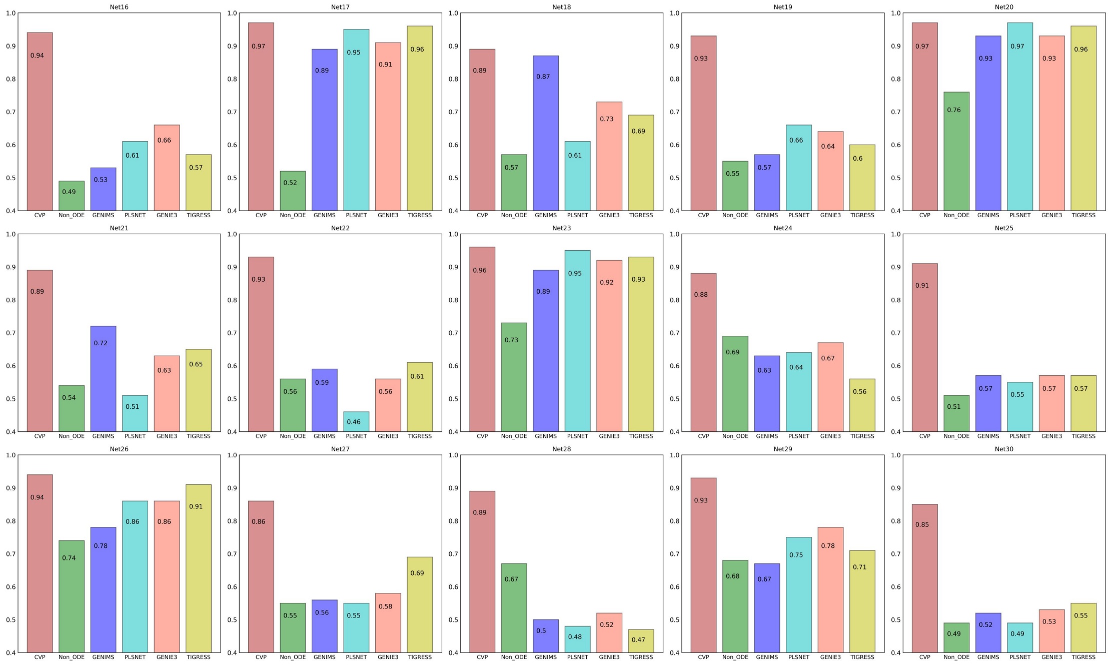


**Fig. S7. The performance of different algorithms on synthesis network of size 100 from DREAM challenges.** The height of the bar chart indicates the accuracy of the different algorithms.

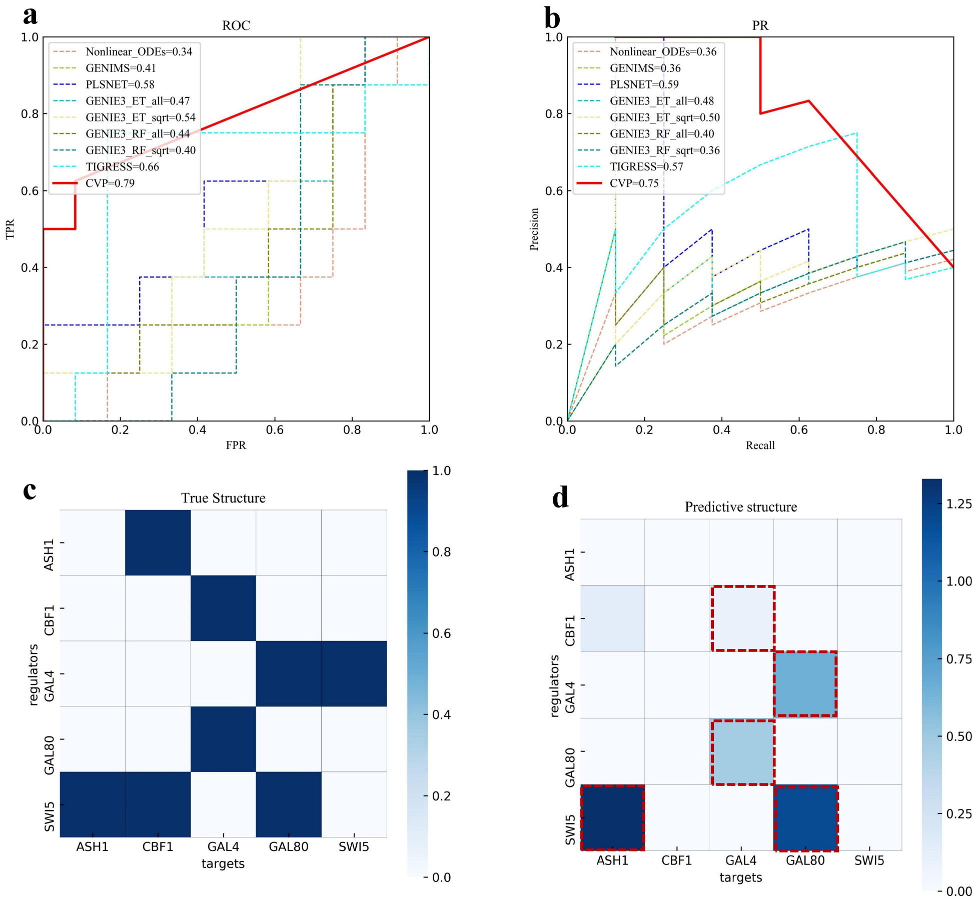


**Fig. S8. The performance of different algorithms on IRMA datasets.** a, the ROC curves of nine methods. b, the PR curves of the nine methods. **c,** heat map of the causal structure of the IRMA data, where dark blue indicates the edges are existent, colorless indicates no edges. d, heat map of the causal structure of the IRMA data detected by the CVP algorithm, with darker colors indicating stronger causal relationships.

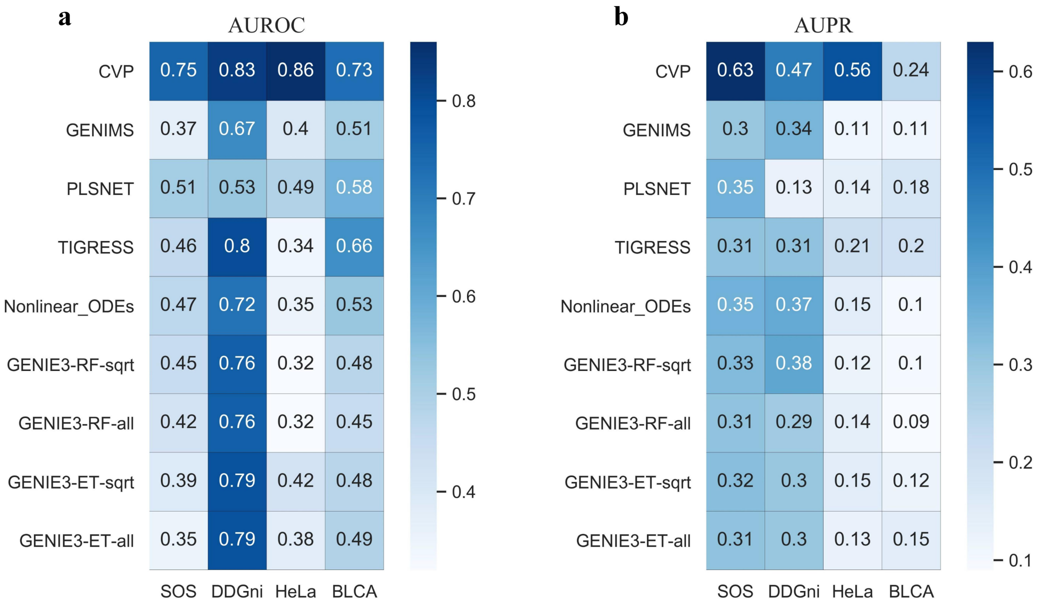


**Fig. S9. Summary of results of different algorithms on real datasets.** a, the AUROC values for the nine algorithms on the four real data sets. b, the AUPR values for the nine algorithms on the four real data sets.

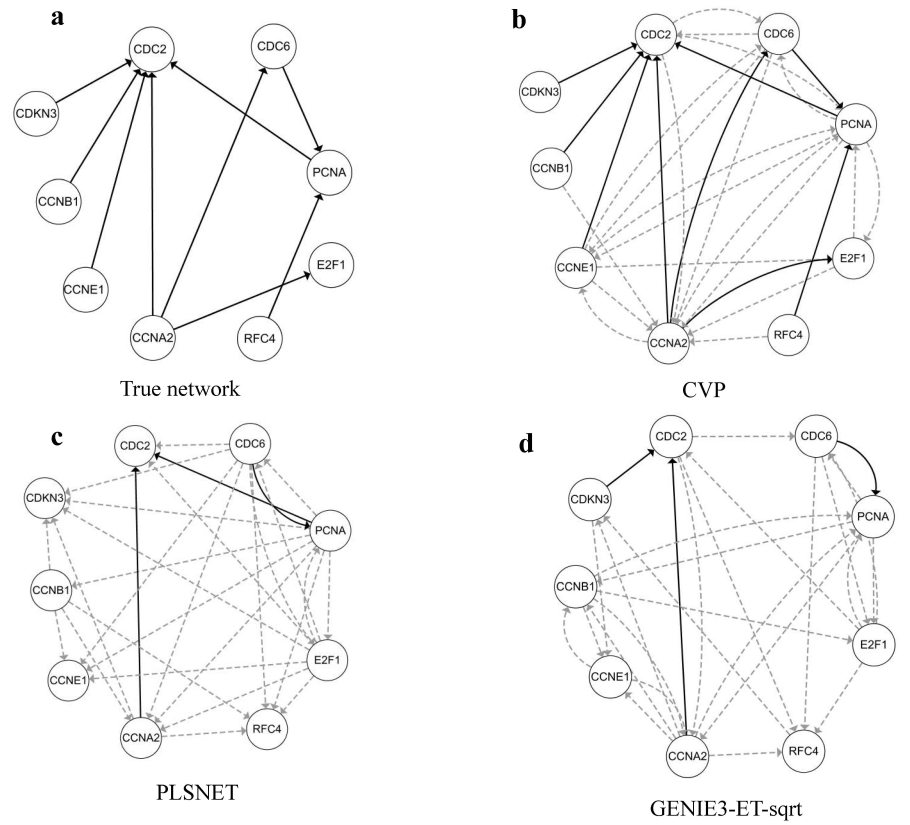


**Fig. S10. The network structure of human HeLa cell cycle dataset.** a, real network of human HeLa cell cycle dataset. b, inferred network by the CVP algorithm. c, inferred network by PLSNET algorithm. d, inferred network by GENIE3-ET-sqrt algorithm.

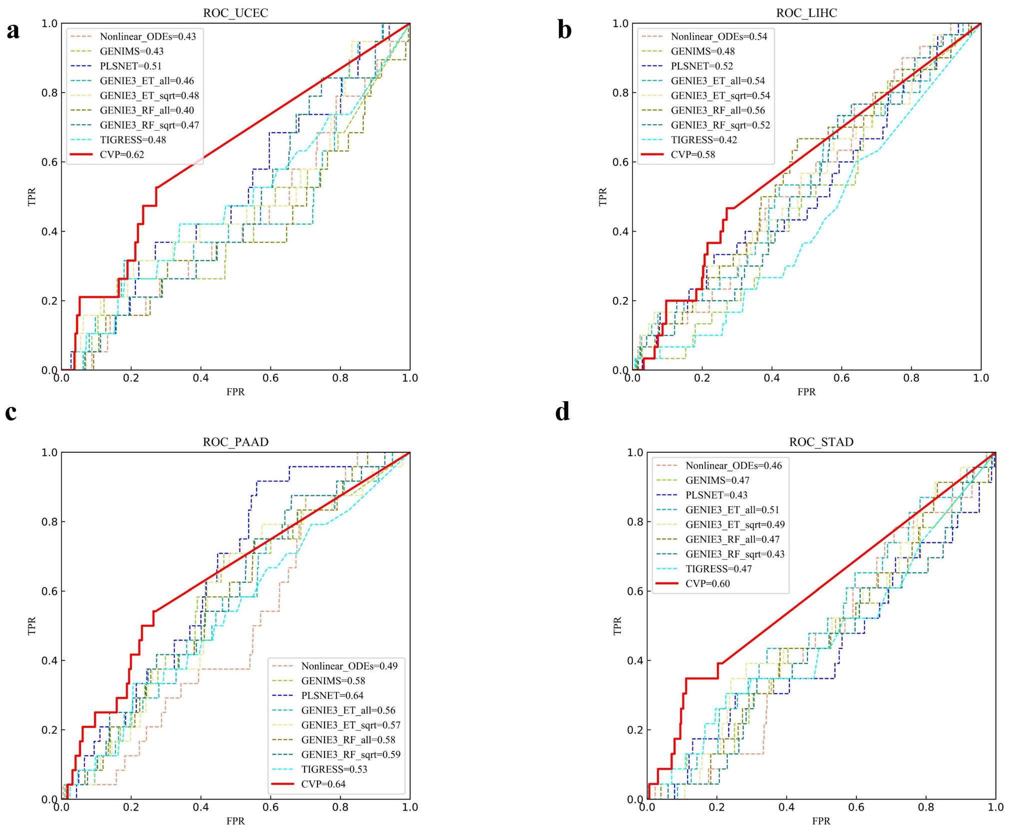


**Fig. S11. Inferred results for the TCGA dataset.** a, b, c and d are the performance of the CVP algorithm and other eight algorithms on the UCEC, LIHC, PAAD and STAD cancer datasets, respectively.

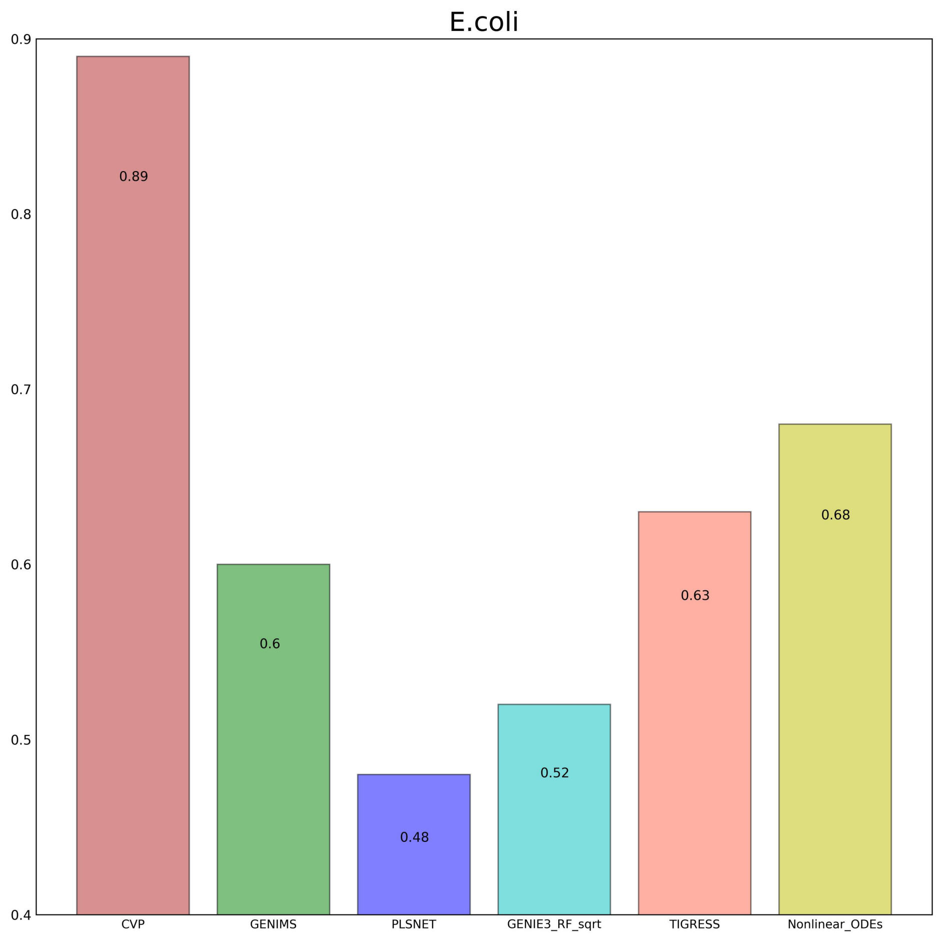


**Fig. S12. Accuracy of six algorithms for large scale networks in *E. coli*.** The horizontal coordinates indicate the name of the algorithm and the vertical coordinates indicate the accuracy rate.

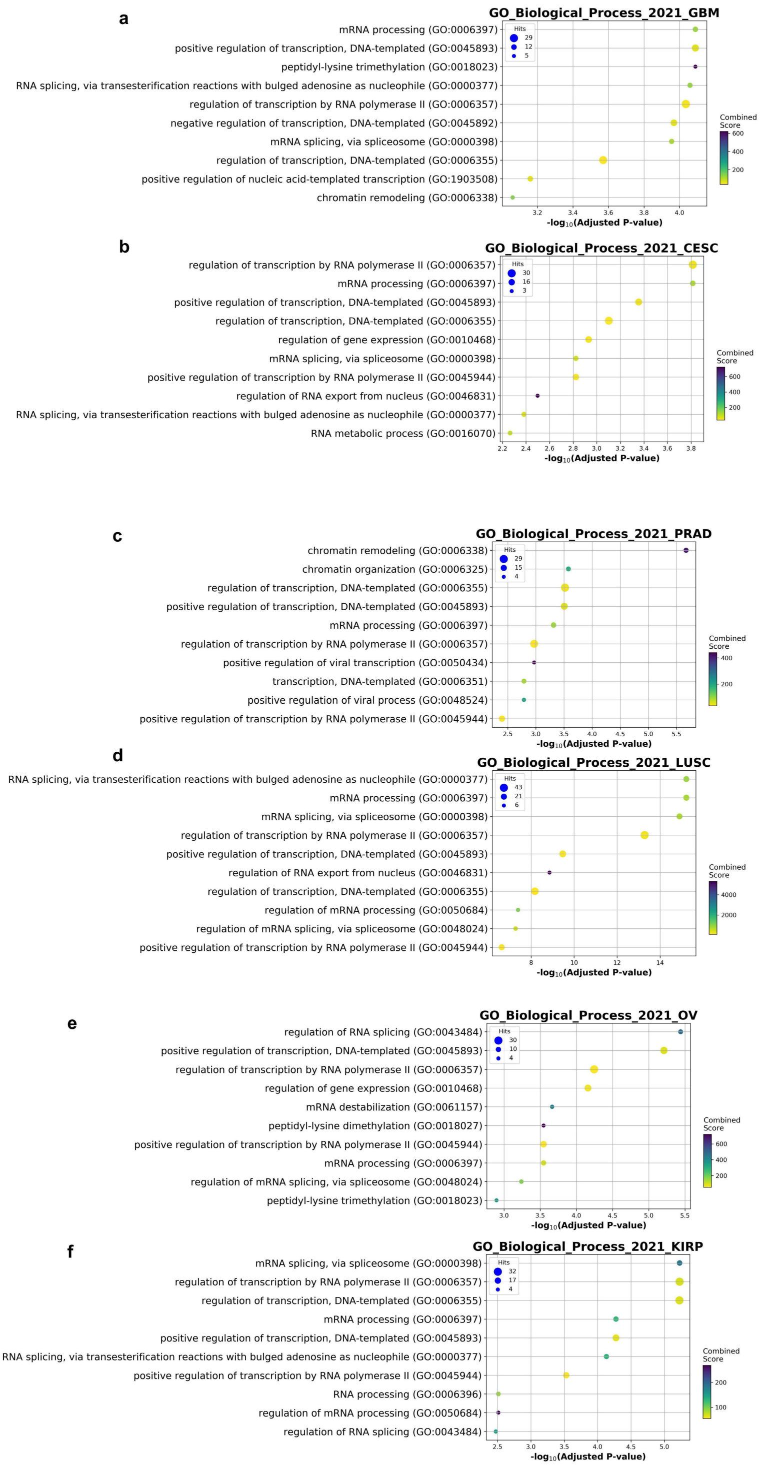


**Fig. S13. The driver genes identified by CVP are involved in important biological functional processes**. a-f are the results of the enrichment analysis of the six cancers in the GO process for GBM, CESC, PRAD, LUSC, OV and KIRP respectively, with darker colors indicating more significant enrichment results.

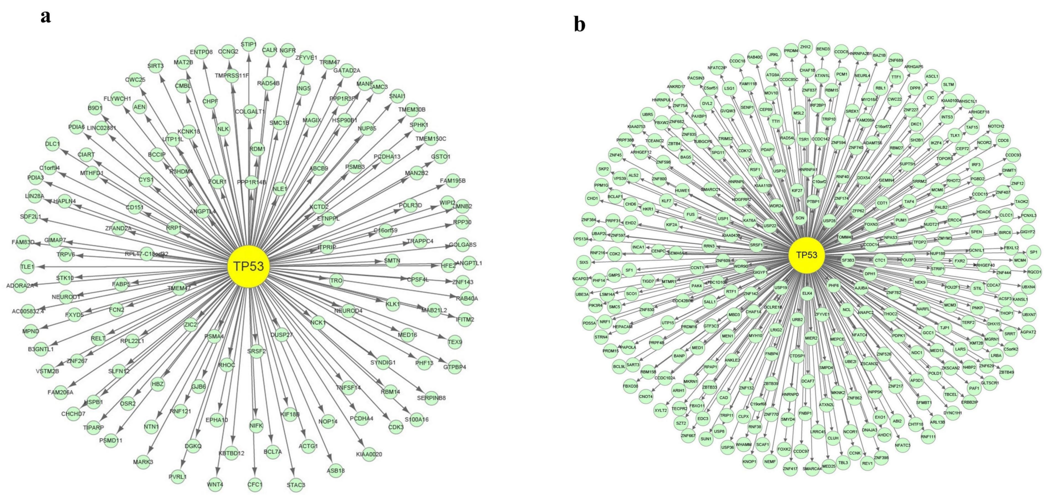


**Fig. S14. The CVP algorithm reveals TP53 regulation patterns in GBM.** a, the regulatory subnetwork of TP53 in the normal data. b, the regulatory subnetwork of TP53 in the GBM cancer data.

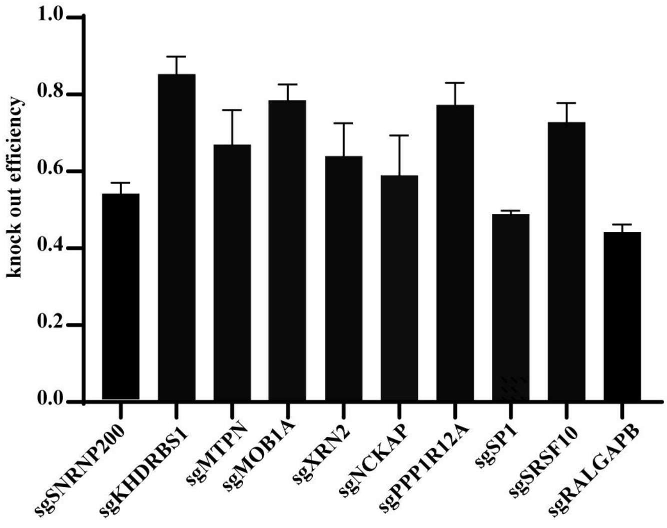


**Fig. S15. The knockdown efficiency of single guide RNAs (sgRNAs) in Huh7 for the top 10 candidate genes.** Huh7 cells were transfected with sgRNAs for 48 h before collecting samples. For each gene knockdown, two sgRNAs were simultaneously transfected. The data are expressed as the gene expression levels relative to the controls tested by qPCR. Error bars represent standard deviations from the mean.

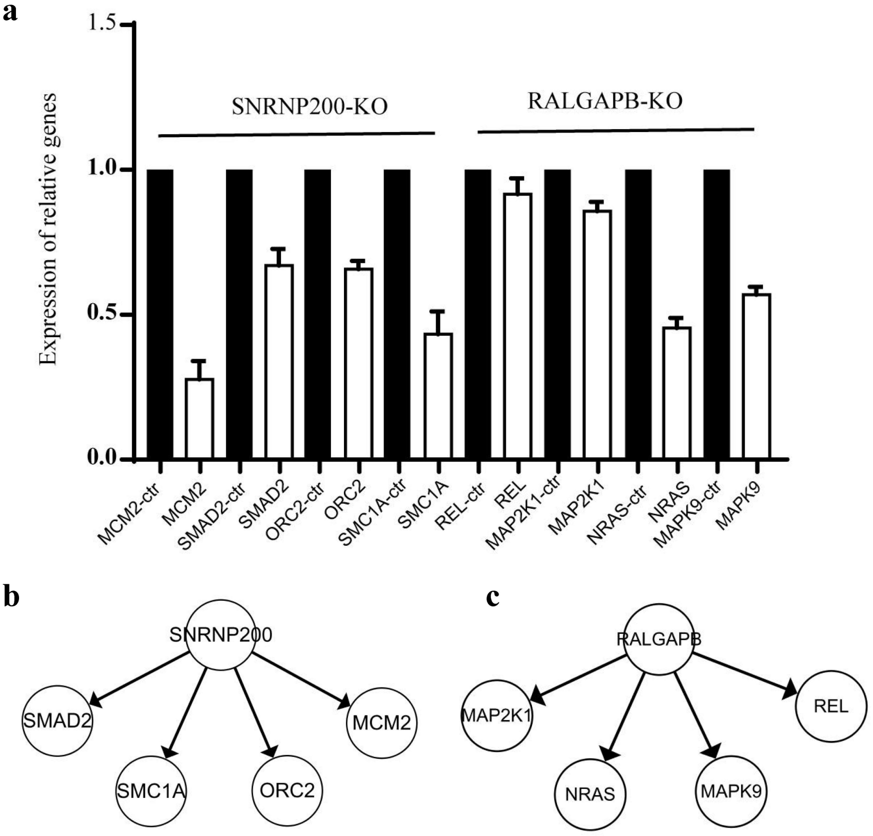


**Fig. S16. The validation of gene regulation in liver cancer cell.** a. The relative expression level of the key inter-regulated genes in Huh7 cells. The experiments were carried out in SNRNP200 and RALGAPB knock out cells. Total RNA was extracted, then cDNA was subjected to real-time PCR. The mRNA expression levels of MCM2, SMAD2, SMC1A, SMC1A, REL, MAP2K1, NRAS and MAPK9 were detected by quantitative Q-PCR. Expression was normalized to that of Actb. Representative results from three independent experiments are shown. ‘ctr’ represents the result in corresponding control group, Mean ± SEM, n = 3 per group. b. The key downstream regulatory genes of SNRNP200. c. The key downstream regulatory genes of RALGAPB.

**Tab. S1.** Generation of causal simulation data for three nodes

| Cases | Directions | Statistical simulation | Statistical distribution | | | |
|---|---|---|---|---|---|---|
| | | | $X$ | $Y$ | $Z$ | $\varepsilon$ |
| Case1 | $X \to Y$ | $Y = 3X + 0.1\varepsilon$ | Normal | | Normal | Uniform |
| Case2 | $X \to Y$<br>$Z \to Y$ | $Y = 3X + 3Z + 0.01\varepsilon$ | Normal | | Normal | Uniform |
| Case3 | $X \to Y$<br>$Y \to Z$ | $Y = 5X + \varepsilon$<br>$Z = Y + \varepsilon$ | Normal | | | Uniform |
| Case4 | $Y \to X$<br>$Y \to Z$ | $X = 5Y + \varepsilon$<br>$Z = 5Y + \varepsilon$ | | Normal | | Uniform |
| Case5 | $X \to Y$<br>$Z \to X$<br>$Z \to Y$ | $X = 3Z + 0.1\varepsilon$<br>$Y = 2X + 5Z + \varepsilon$ | | | Normal | Uniform |

**Tab. S2.** Generation of causal simulation data for eleven nodes

| Cases | Statistical simulation | Statistical distribution | |
|---|---|---|---|
| | | $X_1, X_2, X_3, X_4, X_5, X_6, X_7, X_8, X_9, X_{10}$ | $\varepsilon$ |
| Case1 | $Y = X_1 + X_2 + X_3 + X_4 + X_5 + X_6 + X_7 + X_8 + X_9 + X_{10} + 1 + \varepsilon$ | Uniform | Normal |
| Case2 | $Y = X_2 + X_3 + X_4 + X_5 + X_6 + X_7 + X_8 + X_9 + X_{10} + 1 + \varepsilon$ | Uniform | Normal |
| Case3 | $Y = X_3 + X_4 + X_5 + X_6 + X_7 + X_8 + X_9 + X_{10} + 1 + \varepsilon$ | Uniform | Normal |
| Case4 | $Y = X_4 + X_5 + X_6 + X_7 + X_8 + X_9 + X_{10} + 1 + \varepsilon$ | Uniform | Normal |
| Case5 | $Y = X_5 + X_6 + X_7 + X_8 + X_9 + X_{10} + 1 + \varepsilon$ | Uniform | Normal |
| Case6 | $Y = X_6 + X_7 + X_8 + X_9 + X_{10} + 1 + \varepsilon$ | Uniform | Normal |
| Case7 | $Y = X_7 + X_8 + X_9 + X_{10} + 1 + \varepsilon$ | Uniform | Normal |
| Case8 | $Y = X_8 + X_9 + X_{10} + 1 + \varepsilon$ | Uniform | Normal |
| Case9 | $Y = X_9 + X_{10} + 1 + \varepsilon$ | Uniform | Normal |

**Tab. S3.** Description of the details of the DREAM Challenges data

| Position in thesis | Full name | Data sources | Size |
|---|---|---|---|
| Fig.2a(Network 1) | InSilicoSize10-Yeast3-heterozygous | DREAM3 | 10 |
| Fig. 2b(Network 2) | InSilicoSize10-Yeast2-trajectories | DREAM3 | 10 |
| Fig. 2c(Network 3) | insilico_size10__2_knockdowns | DREAM4 | 10 |
| Fig. 2d(Network 4) | insilico_size10__4_timeseries | DREAM4 | 10 |
| Fig. S5a, S5h | InSilicoSize10-Yeast1-heterozygous | DREAM3 | 10 |
| Fig. S5b, S5i | InSilicoSize10-Yeast3-trajectories | DREAM3 | 10 |
| Fig. S5c, S5j | InSilicoSize10-Yeast1-trajectories | DREAM3 | 10 |
| Fig. S5d, S5k | insilico_size10__3_knockdowns | DREAM4 | 10 |
| Fig. S5e, S5l | insilico_size10__4_knockdowns | DREAM4 | 10 |
| Fig. S5f, S5m | insilico_size10__4_multifactorial | DREAM4 | 10 |
| Fig. S6(Net1) | InSilicoSize50-Ecoli1-heterozygous | DREAM3 | 50 |
| Fig. S6(Net2) | InSilicoSize50-Ecoli1-null-mutants | DREAM3 | 50 |
| Fig. S6(Net3) | InSilicoSize50-Ecoli1-trajectories | DREAM3 | 50 |
| Fig. S6(Net4) | InSilicoSize50-Ecoli2-heterozygous | DREAM3 | 50 |
| Fig. S6(Net5) | InSilicoSize50-Ecoli2-null-mutants | DREAM3 | 50 |
| Fig. S6(Net6) | InSilicoSize50-Ecoli2-trajectories | DREAM3 | 50 |
| Fig. S6(Net7) | InSilicoSize50-Yeast1-heterozygous | DREAM3 | 50 |
| Fig. S6(Net8) | InSilicoSize50-Yeast1-null-mutants | DREAM3 | 50 |
| Fig. S6(Net9) | InSilicoSize50-Yeast1-trajectories | DREAM3 | 50 |
| Fig. S6(Net10) | InSilicoSize50-Yeast2-heterozygous | DREAM3 | 50 |
| Fig. S6(Net11) | InSilicoSize50-Yeast2-null-mutants | DREAM3 | 50 |
| Fig. S6(Net12) | InSilicoSize50-Yeast2-trajectories | DREAM3 | 50 |
| Fig. S6(Net13) | InSilicoSize50-Yeast3-heterozygous | DREAM3 | 50 |
| Fig. S6(Net14) | InSilicoSize50-Yeast3-null-mutants | DREAM3 | 50 |
| Fig. S6(Net15) | InSilicoSize50-Yeast3-trajectories | DREAM3 | 50 |
| Fig. S7(Net16) | InSilicoSize100-Ecoli1-heterozygous | DREAM3 | 100 |
| Fig. S7(Net17) | InSilicoSize100-Ecoli1-null-mutants | DREAM3 | 100 |

| | | | |
|---|---|---|---|
| Fig. S7(Net18) | InSilicoSize100-Ecoli1-trajectories | DREAM3 | 100 |
| Fig. S7(Net19) | InSilicoSize100-Ecoli2-heterozygous | DREAM3 | 100 |
| Fig. S7(Net20) | InSilicoSize100-Ecoli2-null-mutants | DREAM3 | 100 |
| Fig. S7(Net21) | InSilicoSize100-Ecoli2-trajectories | DREAM3 | 100 |
| Fig. S7(Net22) | InSilicoSize100-Yeast1-heterozygous | DREAM3 | 100 |
| Fig. S7(Net23) | InSilicoSize100-Yeast1-null-mutants | DREAM3 | 100 |
| Fig. S7(Net24) | InSilicoSize100-Yeast1-trajectories | DREAM3 | 100 |
| Fig. S7(Net25) | InSilicoSize100-Yeast2-heterozygous | DREAM3 | 100 |
| Fig. S7(Net26) | InSilicoSize100-Yeast2-null-mutants | DREAM3 | 100 |
| Fig. S7(Net27) | InSilicoSize100-Yeast2-trajectories | DREAM3 | 100 |
| Fig. S7(Net28) | InSilicoSize100-Yeast3-heterozygous | DREAM3 | 100 |
| Fig. S7(Net29) | InSilicoSize100-Yeast3-null-mutants | DREAM3 | 100 |
| Fig. S7(Net30) | InSilicoSize100-Yeast3-trajectories | DREAM3 | 100 |

**Tab. S4.** The performance of different methods in IRMA data

| Method | ACC | AUC | PR | TP | FP | TN | FN |
|---|---|---|---|---|---|---|---|
| **CVP** | **0.8** | **0.79** | **0.75** | **5** | **1** | **11** | **3** |
| **GENIMS** | 0.45 | 0.36 | 0.34 | 2 | 5 | 7 | 6 |
| **PLSNET** | 0.6 | 0.58 | 0.59 | **5** | 5 | 7 | **3** |
| **TIGRESS** | 0.7 | 0.66 | 0.57 | 4 | 2 | 10 | 4 |
| **Nonlinear_ODEs** | 0.55 | 0.34 | 0.36 | 2 | 3 | 9 | 6 |
| **GENIE3-RF-sqrt** | 0.45 | 0.4 | 0.36 | 3 | 6 | 6 | 5 |
| **GENIE3-RF-all** | 0.45 | 0.44 | 0.40 | 4 | 7 | 5 | 4 |
| **GENIE3-ET-sqrt** | 0.55 | 0.54 | 0.50 | 4 | 5 | 7 | 4 |
| **GENIE3-ET-all** | 0.55 | 0.47 | 0.40 | 3 | 4 | 8 | 5 |

**Tab. S5.** The performance of different methods in SOS data

| Method | ACC | AUC | PR | TP | FP | TN | FN |
|---|---|---|---|---|---|---|---|
| **CVP** | **0.81** | **0.75** | **0.63** | **13** | **3** | **45** | **11** |
| **GENIMS** | 0.56 | 0.36 | 0.30 | 7 | 15 | 33 | 17 |
| **PLSNET** | 0.60 | 0.51 | 0.35 | 10 | 15 | 33 | 14 |
| **TIGRESS** | 0.625 | 0.46 | 0.31 | 3 | 6 | 42 | 21 |
| **Nonlinear_ODEs** | 0.56 | 0.47 | 0.35 | 7 | 15 | 33 | 17 |
| **GENIE3-RF-sqrt** | 0.64 | 0.45 | 0.33 | 6 | 8 | 40 | 18 |
| **GENIE3-RF-all** | 0.60 | 0.42 | 0.31 | 7 | 12 | 36 | 17 |
| **GENIE3-ET-sqrt** | 0.56 | 0.39 | 0.32 | 5 | 13 | 35 | 19 |
| **GENIE3-ET-all** | 0.64 | 0.35 | 0.31 | 4 | 6 | 42 | 20 |

**Tab. S6.** Results of causal inference for time series data

| Data | DDGni | | HeLa_ | | IRMA | |
|---|---|---|---|---|---|---|
| | AUC | PR | AUC | PR | AUC | PR |
| **CVP** | **0.83** | **0.47** | **0.86** | **0.56** | **0.79** | **0.75** |
| **Nonlinear ODEs** | 0.72 | 0.37 | 0.35 | 0.15 | 0.34 | 0.36 |
| **GENIMS** | 0.67 | 0.34 | 0.32 | 0.1 | 0.36 | 0.34 |
| **PLSNET** | 0.53 | 0.17 | 0.49 | 0.14 | 0.58 | 0.59 |
| **GENIE3_ET_all** | 0.79 | 0.3 | 0.38 | 0.13 | 0.47 | 0.48 |
| **GENIE3_ET_sqrt** | 0.79 | 0.3 | 0.42 | 0.15 | 0.54 | 0.50 |
| **GENIE3_RF_all** | 0.76 | 0.29 | 0.32 | 0.14 | 0.44 | 0.40 |
| **GENIE3_RF_sqrt** | 0.76 | 0.38 | 0.32 | 0.12 | 0.40 | 0.36 |
| **TIGRESS** | 0.8 | 0.31 | 0.34 | 0.21 | 0.66 | 0.57 |
| **GC** | 0.63 | 0.22 | 0.4303 | 0.12 | 0.40 | 0.40 |
| **PTCC+GC** | 0.55 | 0.17 | 0.4303 | 0.12 | 0.65 | 0.50 |

**Tab. S7.** List of driver genes predicted by different algorithms in eight datasets

The table is an Excel file, please access it on below URL:

https://github.com/zyllluck/Table-attachment/blob/main/TabS7.xlsx?raw=true

or

https://sourceforge.net/projects/tabs7/files/TabS7.xlsx/download

**Tab. S8.** The sequences of targeted gRNA expression oligos

| sgRNA | | Guide oligos |
|---|---|---|
| SNRNP200-sgRNA | Sense | ctcgtattgcagactacggg |
| | Antisense | cccgtagtctgcaatacgag |
| | Sense | tagtctgcaatacgagtaca |
| | Antisense | tgtactcgtattgcagacta |
| KHDRBS1-sgRNA | Sense | acgggtcccgacgcgacagt |
| | Antisense | actgtcgcgtcgggacccgt |
| | Sense | gacggcgtctgacgcaccga |
| | Antisense | tcggtgcgtcagacgccgtc |
| MTPN-sgRNA | Sense | ggccctgaaaaacggagact |
| | Antisense | agtctccgtttttcagggcc |
| | Sense | gtgcgacaaggagttcatgt |
| | Antisense | acatgaactccttgtcgca |
| MOB1A-sgRNA | Sense | CTATTCTAAAGCGTCTGTTC |
| | Antisense | GAACAGACGCTTTAGAATAG |
| | Sense | GTTAACTCACCAATCTTAGA |
| | Antisense | TCTAAGATTGGTGAGTTAAC |
| XRN2-sgRNA | Sense | tgacgaacagagacgtgttg |
| | Antisense | caacacgtctctgttcgtca |
| | Sense | ccaacacgtctctgttcgtc |
| | Antisense | gacgaacagagacgtgttgg |
| NCKAP1-sgRNA | Sense | gtccatcatagtcaactgcg |
| | Antisense | cgcagttgactatgatggac |
| | Sense | gagtcccggcgttcttccgc |
| | Antisense | gcggaagaacgccgggactc |
| PPP1R12A-sgRNA | Sense | cgtaattgatgtcggcgccg |
| | Antisense | cggcgccgacatcaattacg |

| | | |
|---|---|---|
| | Sense | gccatcgtcgaacttcacct |
| | Antisense | aggtgaagttcgacgatggc |
| SP1-sgRNA | Sense | CACCGTGGGAAACGCTTCACACGTT |
| | Antisense | AAACAACGTGTGAAGCGTTTCCCA G |
| | Sense | CACCGCGTTTCCCACAGTATGACC |
| | Antisense | AAACGGTCATACTGTGGGAAACGC |
| SRSF10-sgRNA | Sense | ccaacacgtctctgttcgtc |
| | Antisense | gacgaacagagacgtgttgg |
| | Sense | tgacgaacagagacgtgttg |
| | Antisense | caacacgtctctgttcgtca |
| RALGAPB-sgRNA | Sense | ACCTCTCGTCCAACGCTCTC |
| | Antisense | GAGAGCGTTGGACGAGAGGT |
| | Sense | GCAGATTAACGACATACTTC |
| | Antisense | GAAGTATGTCGTTAATCTGC |
| none target | Sense | CTGAGTGAAAAATAAAAGTT |
| | Antisense | GACAATCATGGTGAAAGCGG |
